# Mitochondrial Calcium Signaling Regulates Branched-Chain Amino Acid Catabolism in Fibrolamellar Carcinoma

**DOI:** 10.1101/2024.05.27.596106

**Authors:** Nicole M. Marsh, Melissa J. S. MacEwen, Jane Chea, Heidi L. Kenerson, Albert A. Kwong, Timothy M. Locke, Francisco Javier Miralles, Tanmay Sapre, Natasha Gozali, Madeleine L. Hart, Theo K. Bammler, James W. MacDonald, Lucas B. Sullivan, G. Ekin Atilla-Gokcumen, Shao-En Ong, John D. Scott, Raymond S. Yeung, Yasemin Sancak

**Affiliations:** Department of Pharmacology, University of Washington, Seattle, WA, United States; Department of Surgery, University of Washington Medical Center, Seattle, WA, United States; Department of Chemistry, University at Buffalo, State University of New York, Buffalo, NY, United States; Human Biology Division, Fred Hutchinson Cancer Center, WA, Seattle, United States; Department of Environmental and Occupational Health Sciences, University of Washington, Seattle, WA, United States

**Author notes:** These authors contributed equally (Nicole M. Marsh, Melissa J. S. MacEwen).

## Abstract

Metabolic adaptations in response to changes in energy supply and demand are essential for survival. The mitochondrial calcium uniporter plays a key role in coordinating metabolic homeostasis by regulating TCA cycle activation, mitochondrial fatty acid oxidation, and cellular calcium signaling. However, a comprehensive analysis of uniporter-regulated mitochondrial pathways has remained unexplored. Here, we investigate metabolic consequences of uniporter loss- and gain-of-function using uniporter knockout cells and the liver cancer fibrolamellar carcinoma (FLC), which we demonstrate to have elevated mitochondrial calcium levels. Our results reveal that branched-chain amino acid (BCAA) catabolism and the urea cycle are uniporter-regulated metabolic pathways. Reduced uniporter function boosts expression of BCAA catabolism genes, and the urea cycle enzyme ornithine transcarbamylase (OTC). In contrast, high uniporter activity in FLC suppresses their expression. This suppression is mediated by reduced expression of the transcription factor KLF15, a master regulator of liver metabolism. Thus, uniporter responsive calcium signaling plays a central role in FLC-associated metabolic changes, including hyperammonemia. Our study identifies an important role for mitochondrial calcium signaling in metabolic adaptation through transcriptional regulation of metabolism and elucidates its importance for BCAA and ammonia metabolism in FLC.

## INTRODUCTION

Efficient utilization of energy sources based on their availability and cellular needs requires metabolic flexibility. Mitochondrial calcium (Ca^2+^) signaling plays a central role in metabolic adaptation to acute or chronic changes in energy demands and metabolite levels^1–5^. A principal regulator of this signaling pathway is the mitochondrial Ca^2+^ uniporter (hereon referred to as the uniporter), a Ca^2+^-selective channel in the inner mitochondrial membrane. The uniporter facilitates bulk entry of Ca^2+^ into the mitochondrial matrix^6,7^. This Ca^2+^ influx contributes to activation of the tricarboxylic acid (TCA) cycle, buffering of cytosolic Ca^2+^ signaling, or mitochondrial damage and cell death, depending on the amount of Ca^2+^ that enters the mitochondria^8^. Thus, the uniporter regulates metabolism, mitochondrial and cytosolic Ca^2+^ signaling, and cell survival.

Five core proteins constitute the uniporter: three transmembrane proteins (MCU, EMRE, and MCUb) and two membrane-associated regulatory subunits (MICU1-2)^6,7,9,10^. MCU and EMRE are necessary and sufficient to form a functional Ca^2+^ channel, and MICU1-2 and MCUb regulate uniporter activity^10–13^. In addition, diverse physiological and pathological stimuli alter uniporter-mediated mitochondrial Ca^2+^ uptake through transcriptional and post-translational mechanisms^1,8,14^. Both increased and decreased mitochondrial Ca^2+^ levels are observed in disease-associated states, such as high fat diet-induced obesity^15^,neurodegeneration^16,17^, metabolic disorders^18^, and cancer^19^. Reduced mitochondrial Ca^2+^ uptake leads to slower cell proliferation rates^20–22^, reduced body size^23,24^, exercise intolerance^24^, and epigenetic changes^25^ in mice. Conversely, increased mitochondrial Ca^2+^ load is associated with mitochondrial damage and cell death^26^. Nevertheless, enhanced uniporter activity is thought to be beneficial during energetic stress^1^, pointing to a complex regulation of mitochondrial Ca^2+^ signaling in an effort to balance cellular metabolic needs.

Despite the importance of the uniporter for metabolic adaptation, a comprehensive analysis of mitochondrial pathways that are regulated by the uniporter is lacking. To better understand the effects of the uniporter on mitochondrial function, we analyzed MCU knockout (KO) mitochondria using proteomics and RNA sequencing. Our results show increased expression of branched-chain amino acid (BCAA) catabolism proteins in HeLa MCU KO cells. Furthermore, a key phosphorylation event that inhibits pathway activity is reduced in these cells, thereby augmenting BCAA catabolism.

To determine if these metabolic changes are reversed under conditions of elevated mitochondrial Ca^2+^, we turned to fibrolamellar carcinoma (FLC), an oncocytic tumor with indications of increased mitochondrial Ca^2+^ levels in the literature^27–29^. FLC is a rare liver cancer that primarily affects adolescents and young adults. Recent in-depth analyses of the FLC proteomes and transcriptomes identified substantial differences in tumor mitochondria and metabolism compared to adjacent non-tumor liver samples^30^. These studies indicate a key role for mitochondria in FLC pathogenesis.

Using patient samples and cell models, we show that increased mitochondrial Ca^2+^ levels is a distinguishing feature of FLC. Furthermore, expression of BCAA pathway proteins is attenuated in FLC, a phenotype that is reversed by MCU knockdown. We also report that expression of the transcription factor KLF15, a regulator of multiple BCAA catabolism genes, is regulated by MCU, and is suppressed in FLC tumors. This pathologic event has important implications for FLC patients. KLF15 is vital in promoting expression of the urea cycle enzyme ornithine transcarbamylase (OTC)^31^. Consequently, downregulation of OTC may contribute to impaired urea cycle function and subsequently to hyperammonemia and associated encephalopathy diagnosed in some late-stage FLC patients^32,33^. We observe that, similar to KLF15, OTC expression is suppressed in FLC tumors and is MCU-regulated in cellular models of FLC.

Overall, our results implicate increased uniporter function in FLC and identify BCAA catabolism and the urea cycle as novel uniporter-regulated pathways downstream of KLF15. Based on these findings, we propose that MCU and KLF15 are potential therapeutic targets in the treatment of FLC and its sequelae.

## RESULTS

### Loss of uniporter slows growth and alters expression of mitochondrial proteins

To better characterize how mitochondria respond to loss of mitochondrial Ca^2+^ uptake, we generated a HeLa MCU KO cell line. In parallel we generated a rescue cell line by stable expression of FLAG-tagged MCU on the KO background. Loss of MCU reduced cell growth, which was remedied by MCU re-expression (**Figs. 1A, 1B)**^20–22,34^. Although MCU loss is associated with impaired TCA cycle activity^35^, MCU KO cells did not exhibit a significant decrease in their basal respiration, maximal respiration, ATP production-coupled respiration, respiration due to proton leak, or non-mitochondrial oxygen consumption rates. In addition, they showed only a mild reduction in their spare respiratory capacity (**Figs. S1A-S1G)**.

**Figure 1:**
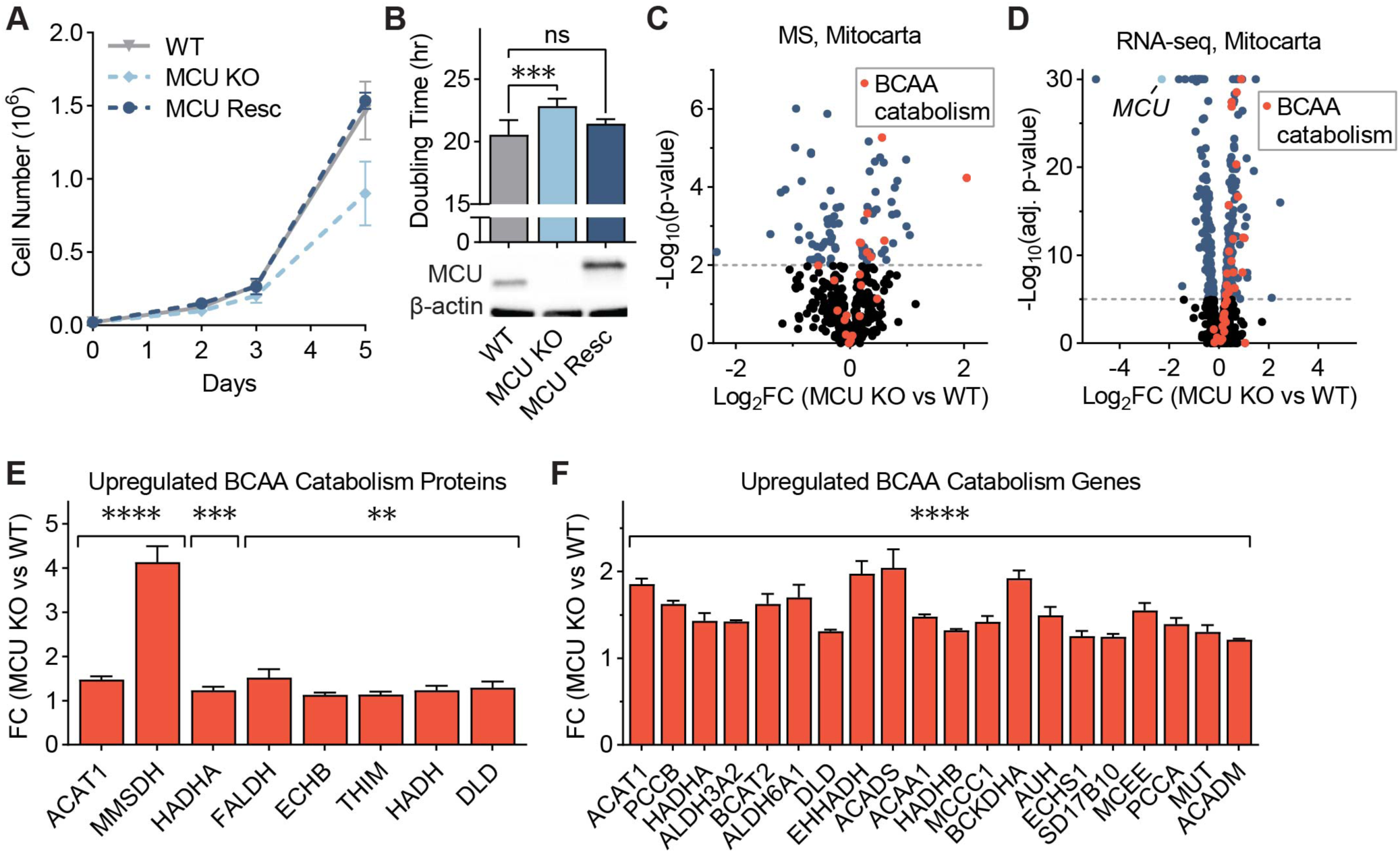
MCU KO cells exhibit growth defects and altered mitochondrial proteome. **(A)** WT, MCU KO, and MCU rescue cells were counted on days 2, 3, and 5 after plating; n=4-6. **(B)** HeLa cell doubling times were calculated from cell counts on days 2 and 5 in (A); statistical significance was determined by Dunnett’s multiple comparisons test following one-way ANOVA; expression of MCU and MCU-FLAG was confirmed by Western blot. **(C)** Volcano plot shows relative abundance of mitochondrial proteins in MCU KO cells compared to WT cells. Red points indicate proteins in the valine, leucine, and isoleucine degradation KEGG pathway; n=5. **(D)** Volcano plot shows relative abundance of mRNAs encoding mitochondrial proteins in MCU KO cells compared to WT cells. Red points indicate genes in the valine, leucine and isoleucine degradation KEGG pathway; MCU is marked in light blue; n=3. **(E, F)** Fold change (FC) of valine, leucine, and isoleucine degradation-associated proteins (E) and genes (F) enriched in MCU KO cells in (C, D); proteins and genes are listed in order of ascending p-value. All error bars represent standard deviation; ns indicates non-significant, * indicates a p-value < 0.05, ** indicates a p-value < 0.01, *** indicates a p-value < 0.001, and **** indicates a p-value < 0.0001.

The lack of a severe oxidative phosphorylation defect in MCU KO cells is attributed to the activation of glycolysis and glutamine utilization^20,34^. The contribution of additional metabolic changes that sustain oxidative phosphorylation and energy production in the absence of the uniporter is unclear. To better characterize the effects of MCU loss on the mitochondria, we compared WT and MCU KO mitochondria using proteomics **(Fig. 1C)** and transcriptomics **(Fig. 1D)**. Gene set enrichment analysis^36^ of significantly upregulated mitochondrial proteins and genes in MCU KO cells showed a strong enrichment of the branched-chain amino acid (BCAA) catabolism pathway and fatty acid oxidation (FAO) pathway in both datasets **(Figs. S1H, S1I).** A link between the FAO and MCU has been reported previously^4^. We confirmed FAO activation in MCU KO cells by lipid analysis. MCU KO cells contained markedly reduced levels of very long-chain fatty acids compared to WT cells **(Fig. S1J)**. This was accompanied by increased levels of acylcarnitines, fatty acid metabolites that are transported to the mitochondria for oxidation **(Fig. S1K)**. Interestingly, 8 of the 48 proteins in the Kyoto Encyclopedia of Genes and Genomes (KEGG) BCAA catabolism pathway (hsa0028) were increased in protein abundance **(Fig. 1E)**. In addition, 20 out of 48 the KEGG BCAA catabolism genes showed 1.3- to 2-fold increase in mRNA levels in MCU KO cells **(Fig. 1F)**. Because of this strong enrichment of BCAA metabolism genes in our dataset and the importance of this process in a diverse set of diseases, we focused on the previously unreported link between mitochondrial calcium signaling and BCAA metabolism.

### BCAA catabolism pathway activity is upregulated by MCU loss

Next, we monitored expression of select pathway proteins after uniporter perturbation. HeLa cells lacking MCU exhibited no mitochondrial calcium uptake, and a small but consistent increase in levels of BCAA catabolism proteins **(Figs. 2A, S2A)**. We observed similar changes in HeLa cells that lack EMRE, an essential component of the uniporter **(Figs. 2B, S2B).** To establish whether uniporter-mediated regulation of BCAA catabolism extends to other species and cell types, in particular to cells that originate from tissues with high BCAA catabolism rates such as liver, we blotted for pathway proteins in the mouse hepatocyte cell line AML12 after RNAi-mediated knockdown (KD) of MCU. MCU KD cells showed reduced mitochondrial calcium uptake rates **(Fig. S2C)** and significantly increased the expression of proteins in the BCAA catabolism pathway **(Fig. 2C).**

**Figure 2.**
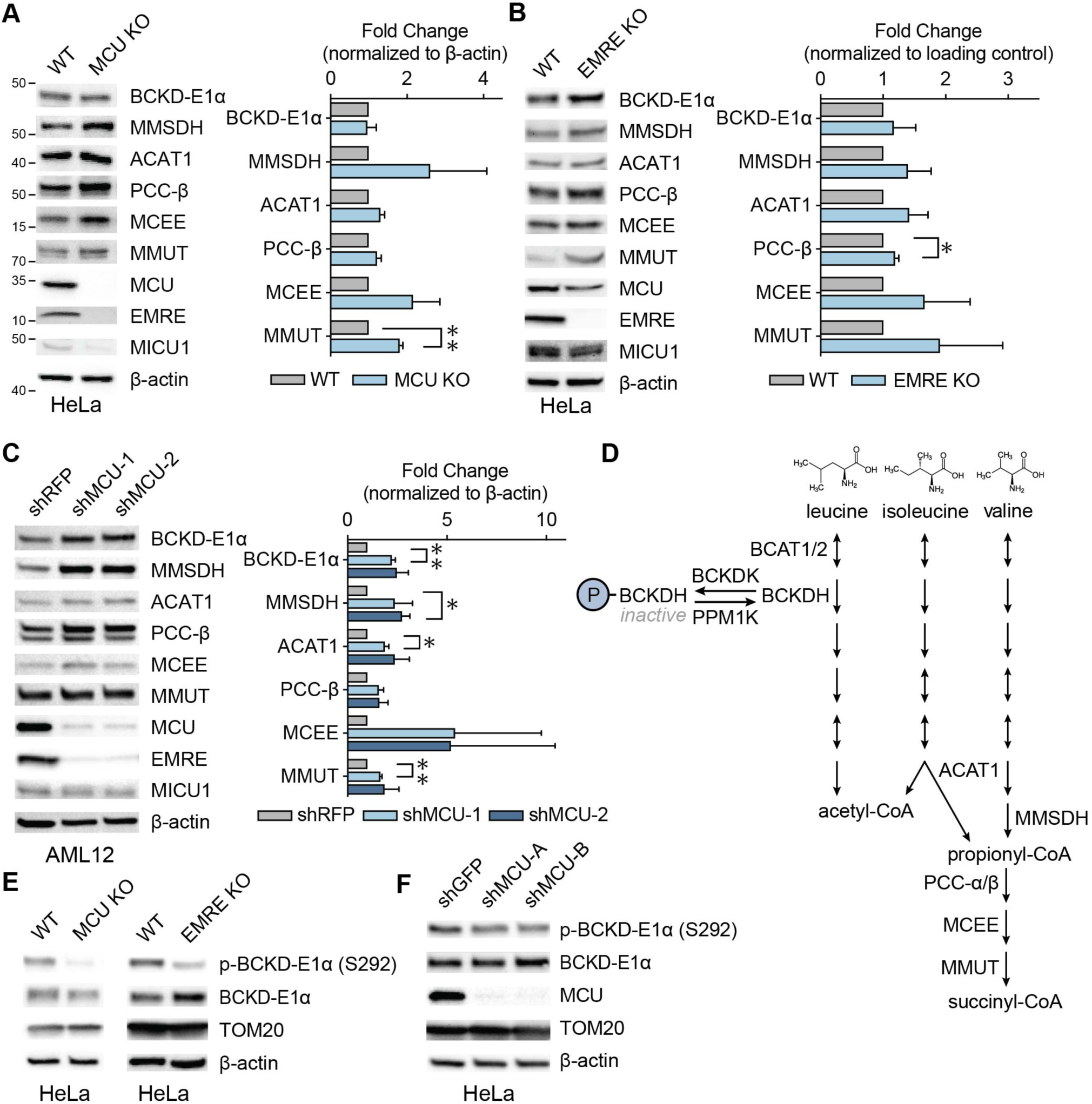
Mitochondrial calcium uniporter regulates BCAA catabolism pathway. **(A)** Representative immunoblots of select BCAA catabolism proteins and uniporter components and their quantification in HeLa WT and MCU KO cells; n=3. **(B)** Representative immunoblots of select BCAA catabolism pathway proteins and their quantification in HeLa WT and EMRE KO cells; n=3. **(C)** Representative immunoblots of select BCAA pathway proteins and their quantification in AML12 cells following shRNA-mediated MCU knockdown; n=3. **(D)** Schematic of BCAA catabolism pathway. The committed step in the pathway is catalyzed by BCKDH complex which is active in the dephosphorylated state in (A-C) was determined by one-sample t-test. All error bars represent standard deviation; * indicates a p-value < 0.05 and ** indicates a p-value < 0.01.

The committed step in the BCAA catabolism pathway is the oxidative decarboxylation reaction carried out by the branched-chain α-ketoacid dehydrogenase (BCKDH) complex. BCKDH activity is regulated by phosphorylation and dephosphorylation of the E1α subunit by the branched-chain α-ketoacid dehydrogenase kinase (BCKDK) and mitochondrial protein phosphatase 1K (PPM1K), respectively. Phosphorylation by BCKDK is inhibitory, whereas dephosphorylation by PPM1K activates the complex **(Fig. 2D)**. We considered the possibility that BCKD-E1α phosphorylation may be uniporter-regulated, in a manner similar to the pyruvate dehydrogenase (PDH) complex in some tissues^37,38^. Interestingly, in HeLa cells, BCKD-E1α phosphorylation was significantly reduced in MCU or EMRE KO cells and after MCU knockdown **(Figs. 2E, 2F, S2D, S2E)**. However, in AML12 cells that originate from liver, where PDH phosphorylation is known to be insensitive to mitochondrial Ca^2+^ levels^37^, we did not observe a change in phospho-BCKD-E1α **(Fig. S2F)**. These data suggest that MCU regulates the activity of the BCAA catabolism pathway both through phosphorylation and transcriptional regulation of pathway enzyme expression in a cell- and tissue-specific manner.

### BCAA catabolism helps maintain NADH/NAD^+^ balance in MCU KO cells

To understand the physiological significance of increased BCAA catabolism in the absence of MCU, we treated cells with the BCKDK inhibitor BT2 and measured cell growth. 10 μM BT2 treatment of WT and MCU KO cells for 3 days reduced BCKD-E1α phosphorylation in both cell lines **(Fig. 3A)**. WT cell growth was not affected, but MCU KO cells showed increased growth **(Fig. 3A)**. In addition to activating BCAA catabolism, BT2 functions as a mild uncoupler^39^. To determine if the increased growth of MCU KO cells with BT2 treatment is due to mild mitochondrial uncoupling, we treated cells with another mild uncoupler 2,4-dinitrophenol (DNP). DNP treatment did not affect cell growth in WT or MCU KO cells **(Fig. S3)**. Therefore, we conclude that activation of BCAA catabolism is beneficial for MCU KO cells. To investigate the mechanism of this unexpected growth phenotype, we first considered the possibility that BCAA catabolism supports the TCA cycle, whose activation is impaired in MCU KO cells^1,24,34^.

**Figure 3:**
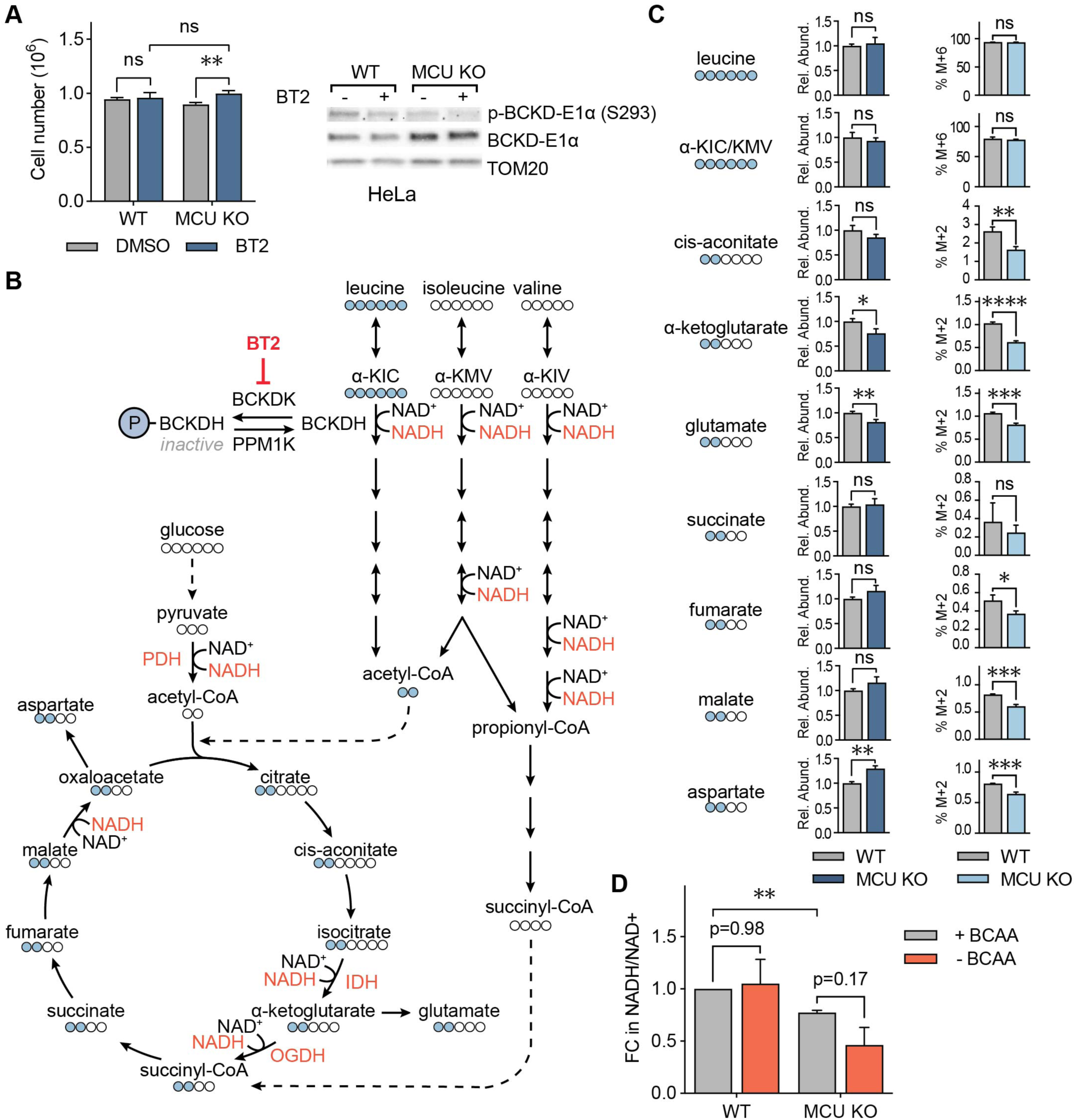
BCAA catabolism maintains NADH/NAD^+^ balance in MCU KO cells. **(A)** Cell numbers of WT and MCU KO cells with and without pharmacological activation of BCAA catabolism by BT2. Cells were counted three days after plating and treatment; n=4-5. Immunoblots show phosphorylated and total BCKD-E1α after vehicle DMSO or BT2 treatment. Statistical significance was determined by Dunnett’s multiple comparisons test following one-way ANOVA. **(B)** Schematic of BCAA catabolism and TCA cycle enzymes and metabolites; BCAA catabolism produces acetyl-CoA and succinyl-CoA, which can enter the TCA cycle; Ca^2+^-regulated enzymes and NAD^+/^NADH coupled reactions are shown in red. Labelled leucine carbons (indicated as blue circles) and their incorporation into the TCA cycle are also shown. **(C)** Relative abundance of indicated metabolites and their fraction generated from labeled leucine in WT and MCU KO HeLa cells. **(D)** Relative NADH/NAD^+^ ratios in WT and MCU KO cells with and without 3hr BCAA starvation are shown. Statistical significance was determined by the Tukey-Kramer test following one-way ANOVA; n=3. All error bars indicate standard deviation; ns indicates non-significant, * indicates a p-value < 0.05. ** indicates a p-value < 0.01, *** indicates a p-value < 0.001, **** indicates a p-value < 0.0001.

Degradation of the three BCAAs – valine, leucine, and isoleucine – generates the TCA cycle metabolites acetyl-CoA and succinyl-CoA **(Fig. 3B)**. Loss of MCU abrogates stimulation of the Ca^2+^-sensitive TCA cycle enzymes pyruvate dehydrogenase (PDH), isocitrate dehydrogenase (IDH), and oxoglutarate dehydrogenase (OGDH). This dampens the production of acetyl-CoA and subsequently succinyl-CoA. Thus, we conjectured that increased BCAA catabolism in MCU KO cells supports the TCA cycle. We tested this hypothesis by tracing ^13^C_6-_labelled leucine carbons after a short labelling period. MCU KO cells had lower levels of total α-ketoglutarate and glutamate **(Fig. 3C)**. Yet, despite having equal amounts of labelled leucine, MCU KO cells showed decreased or unchanged incorporation of leucine-derived carbons into TCA cycle intermediates, as well as into glutamate and aspartate, compared to WT cells **(Fig. 3B, 3C)**. These data suggest that BCAA carbons do not preferentially feed into the TCA cycle in MCU KO cells

As an alternative hypothesis, we speculated that the main role of increased BCAA catabolism in the absence of the uniporter may be to increase NADH production, which is impaired in MCU KO cells^24^. To understand the contribution of the BCAA catabolism pathway to NADH/NAD^+^ homeostasis in MCU KO cells, we starved WT and MCU KO cells of BCAAs for three hours and measured relative NADH/NAD^+^ ratios. As expected, MCU KO cells had a lower NADH/NAD^+^ ratio than WT cells under normal growth conditions^1,34^. BCAA withdrawal did not affect the NADH/NAD^+^ ratio in WT cells, whereas MCU KO cells showed a reduction **(Fig. 3D)**. These results reveal an unexpected function of BCAA catabolism in cellular energy homeostasis through NADH production independent of the TCA cycle activity and define an important role for Ca^2+^ signaling in this regulation.

### Increased mitochondrial Ca^2+^ levels are a hallmark of fibrolamellar carcinoma (FLC)

FLC is a rare liver cancer that affects children and young adults. This cancer is characterized by a somatic ∼400 kb genomic deletion that fuses the first exon of the DnaJ homolog subfamily B member 1 gene (*DNAJB1*) to exon 2–10 of the cAMP-dependent protein kinase catalytic subunit alpha gene (*PRKACA*)^40^ **(Fig. 4A)**. This deletion event generates a fusion kinase, DNAJ-PKAc, which we refer to as DP from hereon. FLC tumors are oncocytic neoplasms that are characterized by an aberrant number of mitochondria^27,28,41^. In addition, earlier reports noted the presence of electron dense particles by electron microscopy (EM) of FLC tumors^28^. These electron dense particles are Ca^2+^ phosphate precipitates that form in the mitochondrial matrix^42,43^ as a result of high matrix Ca^2+^ levels.

**Figure 4.**
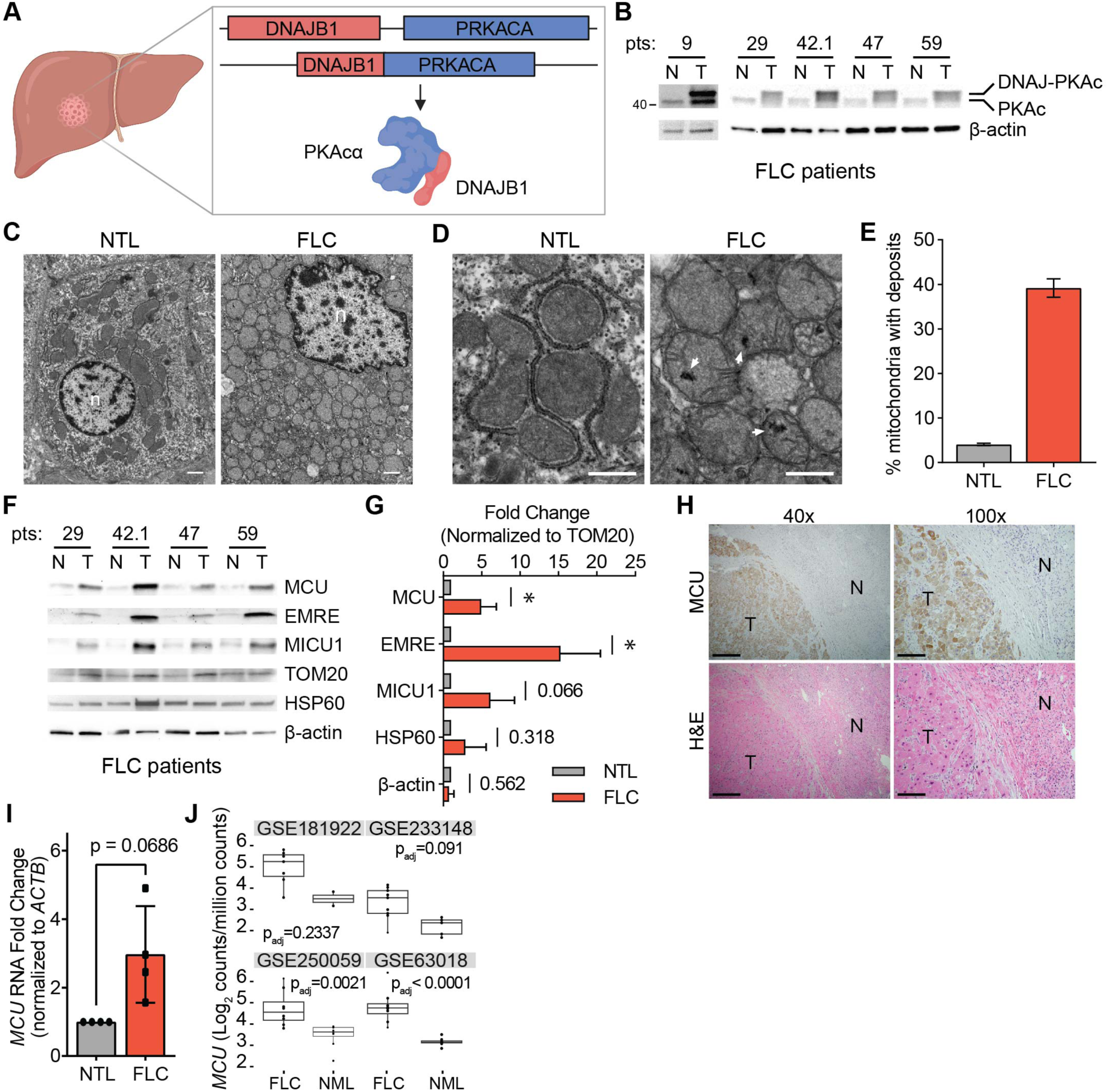
FLC is characterized by increased mitochondrial Ca^2+^ levels and uniporter expression. **(A)** Schematic of FLC liver tumor with heterozygous deletion in chromosome 19 producing the DNAJ-PKAc (DP) fusion protein. **(B)** Immunoblots of lysates from non-tumor (N) and tumor (T) liver from FLC patients show DP fusion protein expression in the tumor. **(C)** Electron micrographs at 10,000x magnification of non-tumor (NTL) and tumor (FLC) sections from patient 9; nuclei are labeled n; scale bars = 1 μm. **(D)** Micrographs of samples shown in (C) at 25,000x magnification; white arrowheads mark representative Ca^2+^ deposits in the tumor; scale bars = 600 nm. **(E)** Percentage of mitochondria from FLC patient 9 with Ca^2+^ deposits; the mean is reported from manual counting of >500 mitochondria per sample by two independent, blinded analysts. **(F)** Immunoblots of uniporter components and control mitochondrial proteins from paired non-tumor (N) and FLC tumor (T) samples. **(G)** Pooled quantification of immunoblots in (F) normalized to TOM20 levels; statistical significance was determined by one-sample t-test. **(H)** H&E and MCU IHC staining of non-tumor (N) and tumor (T) regions of liver from FLC patient 9; 40x and 100x image scale bars are 500 μm and 200 μm, respectively. **(I)** qPCR analysis of MCU RNA expression in paired non-tumor liver (NTL) and tumors from FLC patients 29, 42.1, 47, and 59. **(J)** MCU transcript expression in indicated datasets in tumor (FLC) and normal liver (NML). All error bars indicate standard deviation; numbers above error bars indicate p-values; * indicates a p-value < 0.05.

We sought to further characterize mitochondrial Ca^2+^ changes and investigate uniporter function and BCAA metabolism in FLC. We obtained paired FLC and non-tumor liver (NTL) tissue samples from five patients and confirmed DP expression in the tumors using western blotting with an antibody that detects PKAc **(Fig. 4B, Supp Table 1)**. To investigate mitochondrial Ca^2+^ levels in FLC, we performed EM analysis on tumor and non-tumor liver from one patient **(Figs. 4C, 4D)**. 39% of FLC mitochondria contained one or more electron dense Ca^2+^ deposits compared to 4% of mitochondria from non-tumor liver **(Fig. 4E)**. We also analyzed mitochondria from a second patient, for which a control liver sample did not exist, by comparing mitochondria in oncocytic and peri-oncocytic cells from the tumor periphery **(Figs. S4A-S4C)**. This sample also showed an increased percentage of mitochondria with electron dense particles in oncocytic cells (24%) compared to peri-oncocytic cells (10%) **(Fig. S4D)**. EM analysis of tumor and control liver from a hepatocellular carcinoma (HCC) patient did not show more abundant Ca^2+^ deposits in HCC mitochondria **(Figs. S4E-S4G)**. These observations suggest that high levels of mitochondrial Ca^2+^ deposits may be an FLC-specific phenotype.

We next quantified levels of the uniporter proteins MCU, EMRE and MICU1 in FLC and paired NTL control lysates. MCU and EMRE levels were significantly increased relative to TOM20, and MICU1 trended towards higher expression in tumors **(Figs. 4F, 4G)**. The control mitochondrial protein HSP60 did not show a significant difference between tumors and control lysates, suggesting that the increase in uniporter protein expression is specific and is not simply a result of increased mitochondrial proteome abundance in FLC. Immunohistochemistry (IHC) analysis of tumor and normal liver sections also showed stronger MCU staining in the tumor **(Fig. 4H)**. Quantitative PCR (qPCR) analysis showed that FLC tumors had elevated MCU mRNA levels **(Fig. 4I)**. Analysis of MCU transcript levels in publicly available datasets showed increased MCU expression in FLC tumors compared to normal liver **(Fig. 4J)**. These data are consistent with higher uniporter expression driving increased mitochondrial Ca^2+^ levels in FLC.

### Cellular FLC models show increased mitochondrial Ca^2+^ levels

To better understand the role of mitochondrial Ca^2+^ signaling in FLC metabolism, we turned to two clonal AML12 cell lines, c14 and c4, which were previously generated using genome editing^44^ and used as cellular models of FLC. These clones carry the heterozygous FLC deletion, express a single allele of the fusion kinase DP, and grow faster than WT AML12 cells **(Figs. 5A, 5B)**. The clonal cells also recapitulate FLC-associated signaling events and have been successfully used to investigate FLC biology and therapeutics^44,45^.

**Figure 5.**
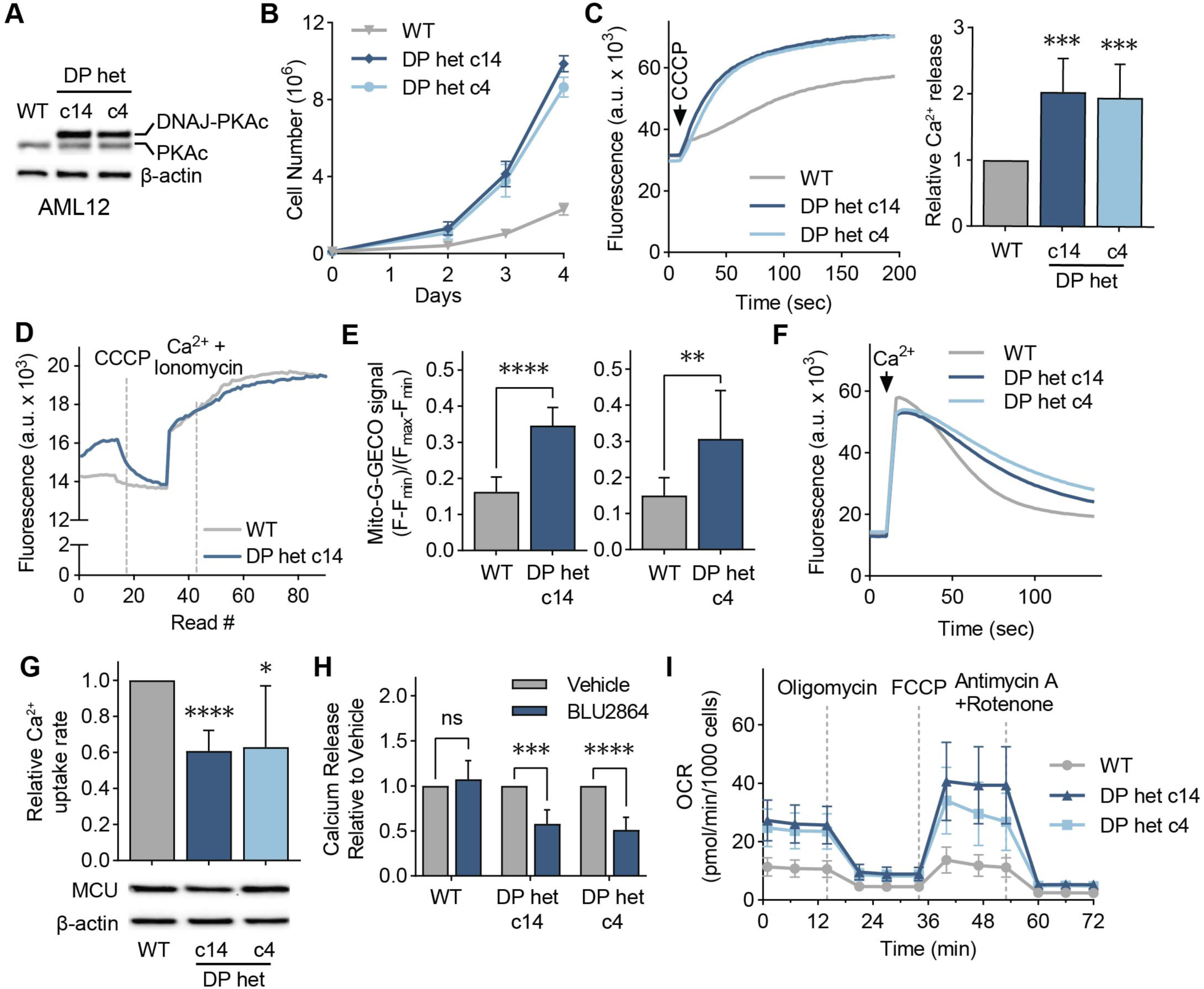
Cellular models of FLC show DP-dependent increase in mitochondrial Ca^2+^ levels. **(A)** Immunoblot of lysates from WT and FLC clones c14 and c4 using an antibody against PKAc. **(B)** Proliferation curves of cellular models of FLC compared to WT AML12 cells; cells were counted on days 2, 3, and 4 after plating; n=3. **(C)** Representative traces and quantification of mitochondrial Ca^2+^ release assays in AML12 cells; cells were treated with the uncoupler CCCP, and the relative amount of Ca^2+^ released was quantified using a Ca^2+^ indicator dye; statistical significance was determined by one-sample t-test; n=10. **(D)** Representative trace of mitochondrial free Ca^2+^ levels of AML12 WT and c14 cells quantified using matrix-targeted Ca^2+^ reporter G-GECO (mito-G-GECO). **(E)** Baseline mito-G-GECO fluorescence normalized to minimum and maximum signals in AML12 WT and c14 or c4 cells; statistical significance was determined by Mann Whitney test; n >12 **(F, G)** Representative traces (F) and mitochondrial Ca^2+^ uptake rates in AML12 cells (G); mitochondrial Ca^2+^ uptake rates were calculated by monitoring Ca^2+^ clearance in the presence of a Ca^2+^ indicator dye; statistical significance was determined by one-sample t-test; n=9. Immunoblot of MCU shows comparable MCU expression in FLC clones compared to WT controls. **(H)** Mitochondrial Ca^2+^ release was assayed as in (C) after cells were treated with 5 μM PKA inhibitor BLU2864 or DMSO for 4 days. Ca^2+^ release was normalized to total protein levels; fold change in released Ca^2+^ is shown relative to DMSO control for each cell line; statistical significance was determined by paired t-test, n=7-9. (I) Seahorse extracellular flux analysis of oxygen consumption rates (OCR) in FLC clones compared to WT AML12 cells at baseline and after indicated treatments; n=10-16. All error bars indicate standard deviation; ns indicates non-significant, * indicates a p-value < 0.05, ** indicates a p-value < 0.01, *** indicates a p-value < 0.001, and **** indicates a p-value < 0.0001.

To determine if c14 and c4 clones have increased mitochondrial Ca^2+^ levels, we treated digitonin-permeabilized cells with the mitochondrial uncoupler CCCP, which results in release of Ca^2+^ ions from mitochondria. Both clones showed increased mitochondrial Ca^2+^ release compared to WT **(Fig. 5C)**. Measurement of free matrix Ca^2+^ levels using the mitochondria-targeted G-GECO Ca^2+^ reporter^46^ indicated higher resting Ca^2+^ levels in the FLC clones **(Figs. 5D, 5E)**. Resting cytosolic Ca^2+^ levels were similar in the clones and WT cells **(Fig. S5A)**. We also measured mitochondrial Ca^2+^ uptake rates in digitonin-permeabilized cells. C14 and c4 cells showed decreased mitochondrial Ca^2+^ uptake rates **(Figs. 5F, 5G),** despite expressing MCU at comparable levels to WT cells **(Fig. 5J)**. This decrease in Ca^2+^ uptake rate was not due to reduced mitochondrial membrane potential in FLC clones **(Fig. S5B)**, and is likely due to higher concentrations of Ca^2+^ in the matrix negatively affecting mitochondrial Ca^2+^ uptake rates. Next, we tested whether elevated mitochondrial Ca^2+^ levels in c14 and c4 cells are dependent on the kinase activity of DP, the fusion protein that sustains FLC tumor growth^47,48^. Interestingly, the PKA inhibitor BLU2864^49^ did not affect the amount of Ca^2+^ released from WT mitochondria, but caused a significant reduction in the FLC clones **(Fig. 5H)**, with concomitant decreases in PKA substrate phosphorylation in all cell lines **(Fig. S5C)**.

To investigate the consequences of increased mitochondrial Ca^2+^ levels for oxidative phosphorylation and energy production in FLC, we measured the oxygen consumption rate (OCR) of AML12 cells with the Seahorse extracellular flux analyzer. FLC clones showed increased basal, maximal, and non-mitochondrial OCR, as well as increased spare respiratory capacity, proton leak- and ATP production-coupled respiration **(Figs. 5I, S5D-S5I)**, as would be expected with increased uniporter function^20,50^. Based on our data, we conclude that high mitochondrial Ca^2+^ is a previously underappreciated feature of FLC that is coupled to oncogenic driver DP activity and alters FLC metabolism.

### Uniporter regulates expression of proteins involved in BCAA catabolism in FLC

Decreased BCAA catabolism is associated with poor patient outcomes in hepatocellular carcinoma^51^. Yet, the mechanisms by which cancer cells downregulate this pathway are not known. We asked whether increased uniporter activity in FLC suppresses expression of BCAA catabolism genes, as opposed to the increase we observed in MCU KO cells. We first analyzed four publicly available FLC gene expression datasets for changes in BCAA catabolism pathway (GO:0009083), focusing on primary tumor and normal liver. 15 out of 22 genes in the pathway showed lower expression in FLC compared to normal liver **(Fig. 6A).** Western blot analysis of tumors from FLC patients also showed significantly reduced levels of BCAA catabolism pathway enzymes compared to matching non-tumor liver samples, with less than 50% expression in the tumors compared to normal liver when protein levels were normalized to the outer mitochondrial membrane protein TOM20 (**Fig. 6B)**. These trends were also evident in FLC clones c14 and c4 **(Fig. 6C).** Knockdown of MCU using two independent shRNAs in FLC cellular models increased expression of BCAA catabolism pathway enzymes **(Fig. 6D)**. Similar to AML12 cells, we did not observe a change in phosphorylation of BCKD-E1α in patient lysates **(Figs. S6A).** This suggests that mitochondrial Ca^2+^ signaling regulates the pathway’s activity through its effects on protein levels in the liver, in contrast to phosphor-regulation observed in HeLa cells. These results are consistent with previous reports^52^, and reflect tissue-specific differences in regulation of mitochondrial metabolism by Ca^2+^ signaling.

**Figure 6:**
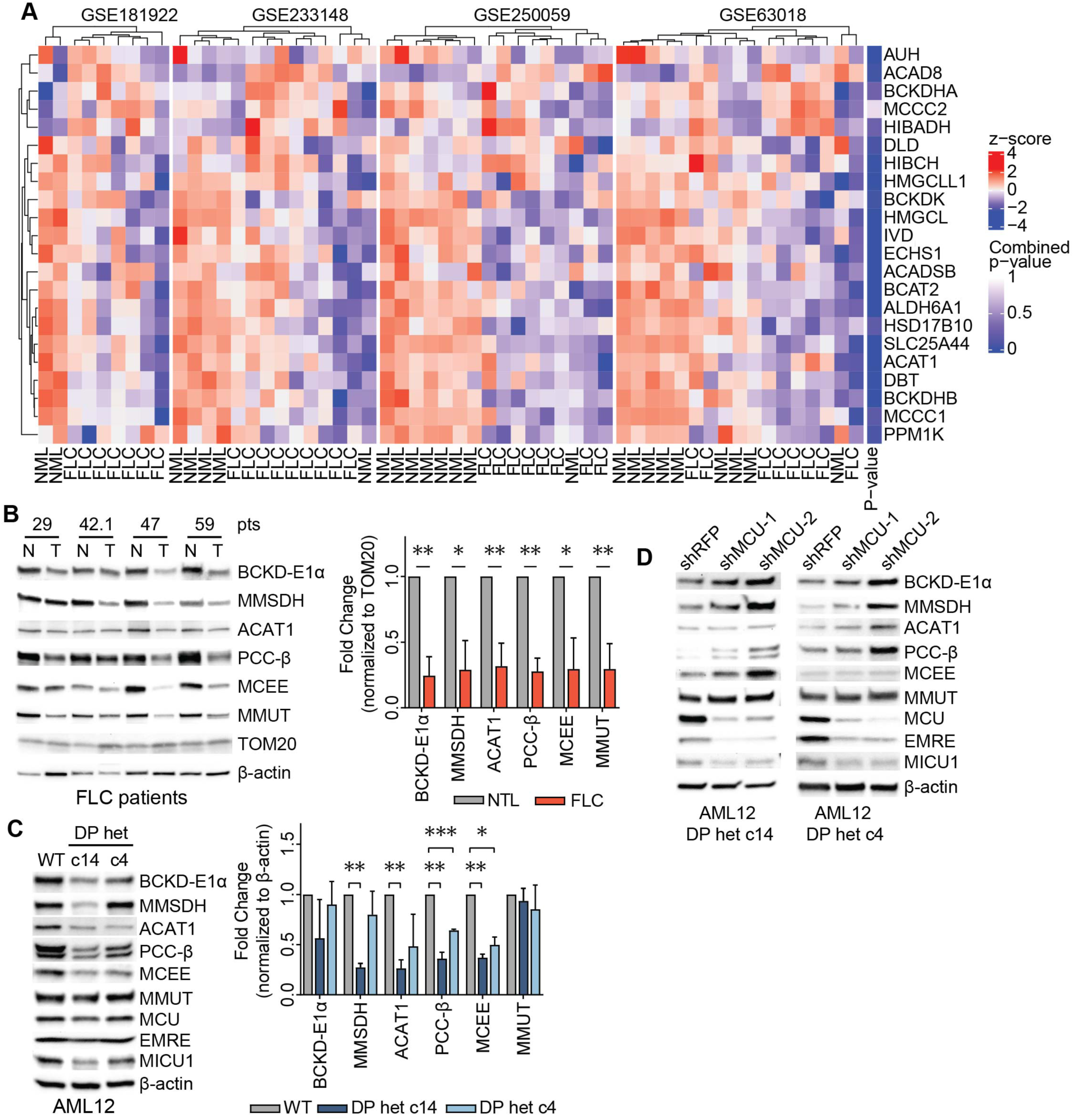
FLC is characterized by uniporter-mediated suppression of BCAA catabolism. **(A)** Analysis of publicly available gene expression datasets of FLC tumor and non-tumor liver (NTL) show decreased in BCAA catabolism gene expression in FLC **(B)** Immunoblots of select BCAA catabolism enzymes from paired non-tumor (N) and FLC tumor (T) lysates and pooled quantification of protein levels normalized to TOM20 levels; statistical significance was determined by one-sample t-test. **(C)** Immunoblots and quantification of select BCAA catabolism proteins and uniporter components in AML12 cells; statistical significance was determined by one-sample t-test; n=4. **(D)** MCU knockdown increases expression of BCAA catabolism pathway proteins in FLC clones; immunoblots of select pathway proteins and uniporter components are shown. All error bars indicate standard deviation; * indicates a p-value < 0.05, ** indicates a p-value < 0.01, and *** indicates a p-value < 0.001.

### Mitochondrial Ca^2+^ signaling regulates expression of KLF15 and its targets in FLC

Reduced mRNA expression of BCAA pathway enzymes in FLC suggested the presence of a transcriptional regulatory mechanism. Krüppel-like factor 15 (KLF15) is a key transcriptional regulator of metabolic gene expression in the liver, including BCAA catabolism and urea cycle genes^31,53,54^. To investigate if KLF15 plays a role in FLC metabolism, we initially probed for KLF15 in lysates from FLC patients by Western blot. FLC tumors exhibited markedly reduced levels of KLF15 **(Fig. 7A)**. Immunohistochemistry of tissue sections from FLC patients confirmed low KLF15 and high MCU expression in the tumors relative to normal liver **(Fig. 7B)**. Likewise, immunoblot analysis demonstrated that KLF15 expression was reduced in AML12 clones c14 and c4 compared to WT cells **(Fig. 7C)**.

**Figure 7:**
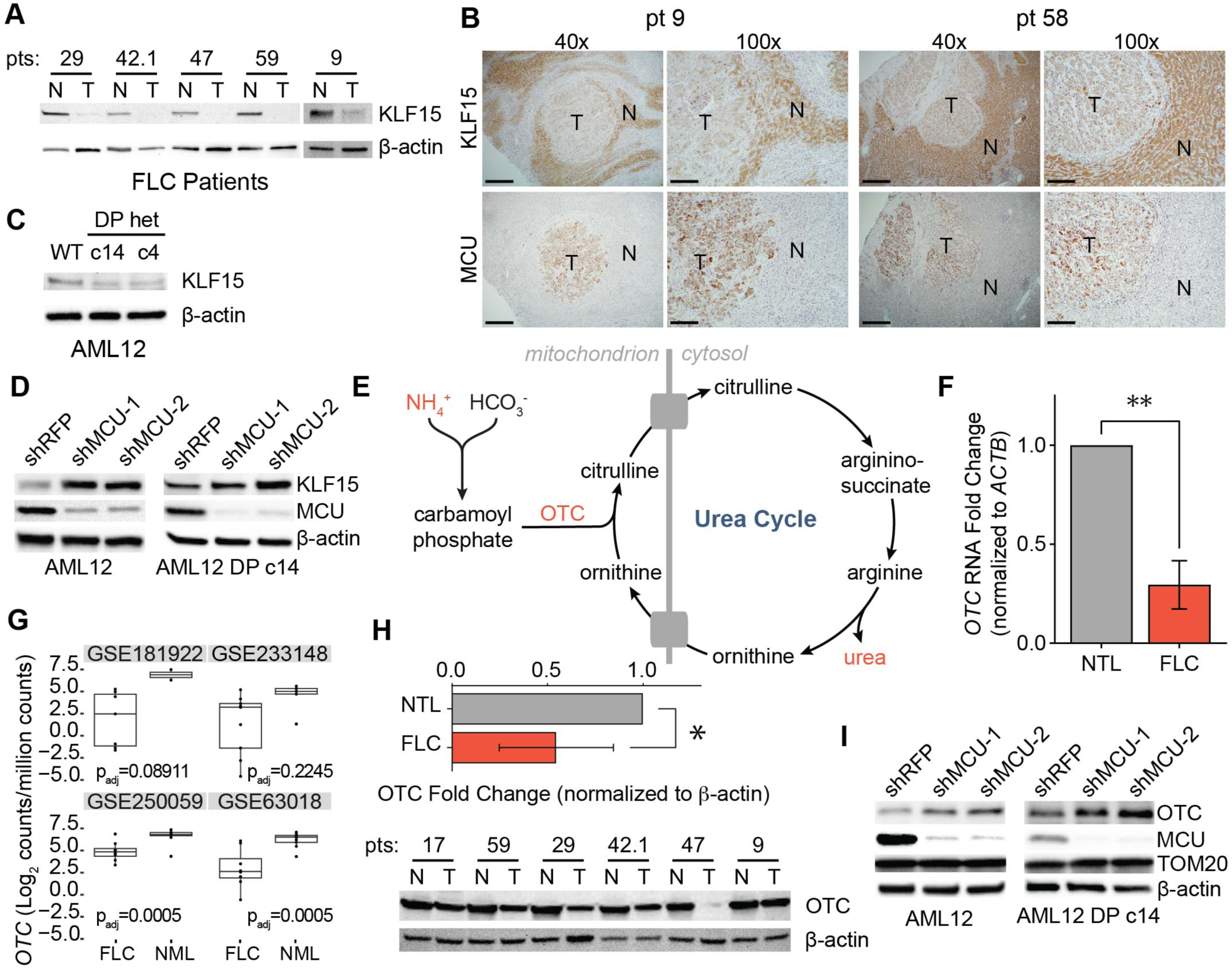
KLF15 and OTC expression are regulated by the uniporter in FLC. **(A)** Immunoblots of KLF15 in paired non-tumor (N) and FLC tumor (T) lysates. **(B)** IHC of MCU or KLF15 on non-tumor (N) and tumor (T) regions from FLC patients 9 and 58; 40x and 100x image scale bars are 500 μm and 200 μm, respectively. **(C)** Immunoblot of KLF15 from WT AML12, c14, and c4 lysates. **(D)** Immunoblots of KLF15 from WT AML12 and c14 lysates following MCU knockdown. **(E)** Schematic of the urea cycle, metabolites, and OTC, a mitochondrial protein with reduced levels in FLC. **(F)** qPCR analysis of *OTC* mRNA expression in paired non-tumor liver and tumors from FLC patients 29, 42.1, 47, and 59; statistical significance was determined by one sample t-test; n=4 **(G)** *OTC* mRNA expression in normal liver (NML) and FLC tumors in the indicated gene expression datasets. **(H)** Immunoblot of OTC and quantification of OTC levels relative to β-actin from paired non-tumor (N) and FLC tumor (T) lysates; statistical significance was determined by one sample t-test; n=5. **(I)** Immunoblots of OTC from WT AML12 and c14 lysates following MCU knockdown. All error bars indicate standard deviation; * indicates a p-value < 0.05 and ** indicates a p-value < 0.01.

To investigate the role of uniporter function in KLF15 regulation, we knocked down MCU in AML12 WT and c14 cells. This caused a substantial increase in KLF15 levels **(Fig. 7D)**, suggesting that mitochondrial Ca^2+^ signaling negatively regulates KLF15 expression. We reasoned that reduced KLF15 expression may have important metabolic consequences beyond inhibition of BCAA catabolism in FLC, including regulation of the urea cycle by KLF15. Defects in the urea cycle can lead to accumulation of ammonia and cause hyperammonemia **(Fig. 7E)**, which can cause neurological damage and death^55^. Decreased OTC expression has also been identified as a potential cause of hyperammonemia in some FLC patients^30^. Interestingly, KLF15 KO mice suffer from increased blood ammonia levels due to significantly reduced OTC expression and activity in the liver^31^. RNA **(Figs. 7F, 7G)** and protein **(Fig. 7H)** levels of OTC showed a significant reduction in tumor samples compared to normal liver in our patient cohort, consistent with transcriptional regulation of OTC in FLC. Moreover, knockdown of MCU in WT AML12 and c14 cells increased OTC expression **(Fig. 7I)**, suggesting a previously unknown link between mitochondrial Ca^2+^ signaling and urea cycle regulation.

Based on our data, we propose a model in which mitochondrial Ca^2+^ signaling regulates the BCAA catabolism pathway and the urea cycle through the transcription factor KLF15 in the liver **(Fig. 8)**. In cells with increased mitochondrial Ca^2+^ levels, such as FLC cells, reduced KLF15 expression leads to reduced expression of BCAA catabolism pathway genes, which conserves essential BCAAs for protein synthesis and promotes cell growth. In cells that do not have bulk mitochondrial Ca^2+^ uptake (MCU-deficient cells), increased KLF15 and activated BCAA catabolism supplies the cells with NADH as a compensatory mechanism in the absence of Ca^2+^ stimulation of TCA enzymes.

**Figure 8:**
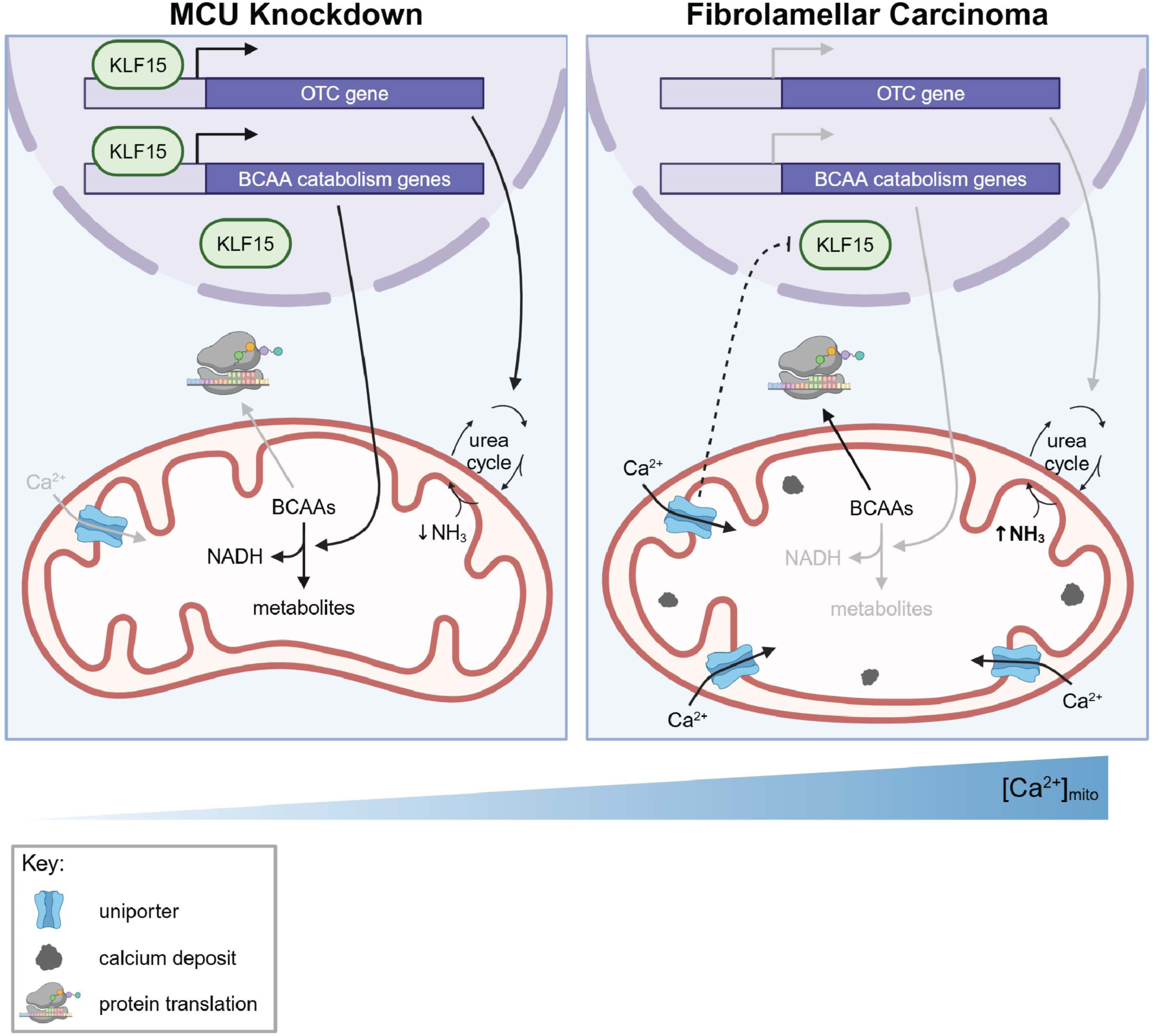
Model for regulation of KLF15, BCAAs, and the urea cycle by mitochondrial Ca^2+^ signaling. Our data suggest that under conditions of low uniporter function in the liver cells, high KLF15 levels stimulate expression of BCAA catabolism pathway genes and OTC. Activation of this pathway helps maintain NADH/NAD^+^ balance. Increased uniporter activity, as observed in FLC, inhibits KLF15, leading to decreased BCAA catabolism enzyme and OTC expression. This inhibition conserves BCAAs for translation and cell growth. MCU inhibition also causes urea cycle impairment, which can lead to hyperammonemia. How the uniporter regulates KLF15 expression is not known.

## DISCUSSION

The BCAA catabolism pathway is responsive to nutrients, hormones, and circadian rhythm. Flux though this pathway is altered in disease states including type II diabetes, cardiovascular disease, liver disease, and cancer^56–58^. Here, we identify mitochondrial Ca^2+^ signaling as a previously unrecognized mechanism by which the BCAA catabolism pathway is regulated. More specifically, we demonstrate that loss of uniporter function results in increased expression of BCAA catabolism enzymes and activation of the pathway’s committed step. Conversely, increased mitochondrial Ca^2+^ uptake inhibits BCAA catabolism transcriptionally. Based on our findings, we propose that mitochondrial Ca^2+^ signaling acts as a regulatory switch through which BCAAs are either broken down to support cellular NADH/NAD^+^ balance, or conserved for translation in accordance with the cell’s needs **(Fig. 8)**.

Our findings have additional implications for understanding FLC pathology. First, we show that increased mitochondrial calcium levels are linked to kinase activity of the oncogene DP in this cancer. Second, reduced expression of the KLF15 target OTC is thought to contribute to hyperammonemia observed in FLC patients^30,33^. Moreover, accumulation of the OTC substrates ammonia and ornithine can support the production of collagen^59–61^, which is a large component of the fibrous bands that are hallmarks of FLC tumors. Interestingly, elevated levels of MCU, and downregulation of KLF15 are associated with increased collagen production and fibrosis various tissues, supporting a role for this pathway in FLC fibrosis^62–65^. Although the mechanism of DP-induced increase in mitochondrial Ca^2+^ signaling remains to be investigated, we delineate a pathway that starts with DP-induced activation of mitochondrial Ca^2+^ signaling, and subsequent KLF15-dependent transcriptional changes, urea cycle defects, and possibly fibrosis in FLC.

Our results indicate that the transcription factor KLF15, a known regulator of BCAA catabolism enzymes^54,66^, mediates the effects of MCU on BCAA pathway gene expression. This suggests that KLF15 may play a broader role in transcriptional regulation of metabolism by mitochondrial Ca^2+^ signaling, especially in the context of cancer. Importantly, reduction in KLF15 levels has been correlated with tumor growth and poor prognosis in several types of cancer^67–72^. However, to our knowledge, this is the first time KLF15 expression changes have been implicated in liver cancer. The MCU/KLF15/BCAA catabolism signaling axis that we identified in FLC may also play a role in other forms of cancer. Based on analysis of the Cancer Genome Atlas (TGCA), the BCAA catabolism pathway is downregulated in approximately 70% of cancers^51^. Additionally, both suppressed BCAA catabolism and increased uniporter activity correlate with poor patient prognosis in hepatic cancers^51,73–75^. This raises the intriguing possibility that uniporter-mediated metabolic changes may be a more general mechanism of metabolic regulation in cancer, whereby increased mitochondrial Ca^2+^ levels stimulate TCA cycle activity, downregulate BCAA catabolism, and conserve essential amino acids for protein synthesis and cell proliferation.

The majority of BCAA catabolism occurs in skeletal muscle, adipose tissue, and the liver^65^. The initial conversion of BCAAs to α-ketoacids is catalyzed by cytosolic BCAT1 and mitochondrial BCAT2 enzyme. Recent findings suggest that human and murine liver show differences in liver BCAT activity^76,77^. BCAT activity is present in the human liver, whereas murine hepatocytes, including the AML12 model we used in this study, lack BCAT activity. As such, we were unable to measure changes in BCAA consumption in these samples. This limited our ability to assess rates of BCAA catabolism directly. Nonetheless, regulation of the pathway components remains sensitive to alterations in cell lines, enabling us to characterize upstream effectors regardless of pathway activity *in vitro*.

PKA signaling regulates many aspects of cancer biology^78^. Our work presents a new, mitochondrial Ca^2+^ signaling-dependent mechanism downstream of PKA signaling on regulation of cancer metabolism. Although it how DP expression increases MCU activity remains to be identified, our work suggests that MCU and KLF15 are potential therapeutic targets in the treatment of FLC and its sequelae.

## MATERIALS and METHODS

### Cell lines and tissue culture

All cell lines were grown in a standard tissue culture incubator (VWR, 75875-212) at 37°C with 5% CO_2._ HeLa cells were cultured in DMEM (Thermo Fisher Scientific, 11-965-118) supplemented with 10% FBS (VWR, 89510-186), 2 mM GlutaMAX (Thermo Fisher Scientific, 35-050-061), and 100 IU/mL penicillin and 100 μg/mL streptomycin (VWR, 45000-652). HeLa cells were obtained from the Whitehead Institute for Biomedical Research. The HeLa cell line has the following short tandem repeat profile: D5S818 (11, 12); D13S317 (12, 13.3); D7S820 (8, 12); D16S539 (9, 10); vWA (16, 18); TH01 (7); AMEL (X); TPOX (8, 12); CSF1PO (9, 10). This profile is a 94% match to HeLa Cervical Adenocarcinoma (Human) (CCL-2; ATCC) according to the analysis performed by the ATCC database. AML12 cells were cultured in DMEM/F-12 medium (Gibco, 11330057) supplemented with 10% FBS (VWR, 89510-186), 1X insulin-transferrin-selenium (Gibco, 41400045), 40 ng/mL dexamethasone (MP Biomedicals, 0219456180), and 100 IU/mL penicillin and 100 μg/mL streptomycin (VWR, 45000-652). AML12 cells were obtained from ATCC (CRL-2254, lot #70039497) and CRISPR-edited AML12 DNAJ-PKAc clones c14 and c4 were generated and validated by the Scott Lab as described^44^. All cell lines were tested for mycoplasma using the Genlantis MycoScope PCR Detection Kit (VWR, 10497-508) and were confirmed to be free of mycoplasma contamination.

### Generation of knockout and rescue cell lines

Broad Institute gRNA sequences for *MCU* (TGAACTGACAGCGTTCACGC) and *EMRE* (GTCTCAGCCAGGTACCGTCG) were each cloned into the pU6T7 vector (Addgene, 71462). 5 million HeLa cells were resuspended in 400 μL DMEM with 2 μg SpCas9 plasmid (Addgene, 71814), 500 ng gRNA plasmid, and 17.5 μg pUC19 plasmid (Addgene, 50005). The cells and DNA were transferred to a cuvette, incubated on ice for 5 min, and electroporated at 200 V with a capacitance of 950 μF and pulse length of 23 msec. Electroporated cells were allowed to grow to confluence on a 10 cm plate before being diluted to 1 cell/200 μL and plated in 96-well plates to obtain single cell clones. HeLa MCU KO and EMRE KO clones were verified by Western blotting, functional assays, and sequencing. To rescue MCU expression, MCU KO clone 18 was infected with lentivirus containing an MCU-FLAG construct with the sequence below. Following resistance marker selection, single cell clones were isolated and screened for MCU expression by Western blot.

### MCU-FLAG DNA sequence

ATGGCGGCCGCCGCAGGTAGATCGCTCCTGCTGCTCCTCTCCTCTCGGGGCGGCGGCGGCGGGGGCGCCGGCGGCTGCGGGGCGCTGACTGCCGGCTGCTTCCCTGGGCTGGGCGTCAGCCGCCACCGGCAGCAGCAGCACCACCGGACGGTACACCAGAGGATCGCTTCCTGGCAGAATTTGGGAGCTGTTTATTGCAGCACTGTTGTGCCCTCTGATGATGTTACAGTGGTTTATCAAAATGGGTTACCTGTGATATCTGTGAGGCTACCATCCCGGCGTGAACGCTGTCAGTTCACACTCAAGCCTATCTCTGACTCTGTTGGTGTATTTTTACGACAACTGCAAGAAGAGGATCGGGGAATTGACAGAGTTGCTATCTATTCACCAGATGGTGTTCGCGTTGCTGCTTCAACAGGAATAGACCTCCTCCTCCTTGATGACTTTAAGCTGGTCATTAATGACTTAACATACCACGTACGACCACCAAAAAGAGACCTCTTAAGTCATGAAAATGCAGCAACGCTGAATGATGTAAAGACATTGGTCCAGCAACTATACACCACACTGTGCATTGAGCAGCACCAGTTAAACAAGGAAAGGGAGCTTATTGAAAGACTAGAGGATCTCAAAGAGCAGCTGGCTCCCCTGGAAAAGGTACGAATTGAGATTAGCAGAAAAGCTGAGAAGAGGACCACTTTGGTGCTATGGGGTGGCCTTGCCTACATGGCCACACAGTTTGGCATTTTGGCCCGGCTTACCTGGTGGGAATATTCCTGGGACATCATGGAGCCAGTAACATACTTCATCACTTATGGAAGTGCCATGGCAATGTATGCATATTTTGTAATGACACGCCAGGAATATGTTTATCCAGAAGCCAGAGACAGACAATACTTACTATTTTTCCATAAAGGAGCCAAAAAGTCACGTTTTGACCTAGAGAAATACAATCAACTCAAGGATGCAATTGCTCAGGCAG AAATGGACCTTAAGAGACTGAGAGACCCATTACAAGTACATCTGCCTCTCCGACAAATTGGTGAAAAAGATTCTAGAGGTGGATCTGGTGGATCTGGTGGATCTATGGATTACAAGGATGACGATGACAAG

### Proliferation assays

20,000 HeLa cells were plated in 6-well plates on day 0. On days 2, 3, and 5, the media was aspirated, and the cells were washed with PBS (Thermo Fisher Scientific, 20012050), detached with 200 μL trypsin solution (Gibco, 12605-010), and then resuspended in 1.8 mL media for a total volume of 2 mL. The resuspended cells were counted using a Beckman Coulter Z2 Cell and Particle Counter (Beckman Coulter, 383550). 100,000 AML12 cells were plated in 10 cm plates on day 0. On days 2, 3, and 4, the media was aspirated, and the cells were washed with PBS (Thermo Fisher Scientific, 20012050), detached with 2 mL trypsin solution (Gibco, 12605-010), and then resuspended in 4 mL media, for a total volume of 6 mL. The resuspended cells were counted using a Coulter Z2 Cell and Particle Counter (Beckman Coulter, 383550).

### Mass spectrometry sample preparation

Five replicates of 500,000 WT and MCU KO HeLa cells were plated in 10 cm plates. After 48 hr, the media was aspirated, and cells were washed with cold PBS (Thermo Fisher Scientific, 20012050) and lysed with 350 μL fresh lysis buffer consisting of 8 M urea (Sigma-Aldrich, U5128), 1 M Tris, 1 M tris(2-carboxyethyl)phosphine hydrochloride (TCEP, Sigma-Aldrich, C4706), and 0.6 M 2-chloroacetamide (CAM, Sigma-Aldrich, C0267). Lysates were sonicated on ice for 10 min with 30 sec on/off pulses at an amplitude setting of 100 in a cup-horn sonicator. Samples were vortexed at 1400 rpm on a thermomixer (Eppendorf) for 30 min at 37°C. Samples were then centrifuged at 17,000 g for 10 min at 4°C, and 175 μL of the supernatant was transferred to a new tube. 700 μL cold acetone was added to each sample. 43.75 μL trichloroacetic acid (TCA, Sigma-Aldrich, T4885) was then added for a final concentration of 5%. The samples were vortexed briefly at 1400 rpm and stored overnight at -20°C. Samples were centrifuged at 6,000 g for 10 min at 4°C, and the supernatant was aspirated. 500 μL cold acetone was added to the pellet, and the pellet was sonicated in an ice bath until the pellet was dissolved (5-10 min). The samples were incubated on ice for another 15 min then centrifuged at 6,000 g for 10 min at 4°C. The supernatant was aspirated, and the pellets were air dried for 2-3 min. Samples were resuspended in 200 μL 8 M urea, 100 mM Tris solution. Samples were vortexed for 1 hr at 37°C to dissolve the pellet in solution. Protein concentrations were measured with the Pierce 660 nm Protein Assay (Thermo Fisher Scientific, 22660) and 500 μg of protein per sample was used for subsequent mass spectrometry experiments. Samples were diluted 1:1 with 100 mM triethylammonium bicarbonate (TEAB, Sigma, T7408), reducing the urea concentration to 4 M. A stock of 2 μg/μl Lys-C endoproteinase (Fujifilm Wako Chemicals, 121-05063) was added to each sample for an enzyme:protein ratio of 1:100. Samples were vortexed at 1400 rpm for 2 hr at 37°C. Samples were again diluted 1:1 with 100 mM TEAB, bringing the urea concentration to 2 M. 0.5 μg/μl stock of trypsin (Promega, V5111) dissolved in 40 mM acetic acid was added to each sample for an enzyme:protein ratio of 1:100. Samples were vortexed at 1400 rpm overnight at 37°C. The following day, HPLC-grade formic acid (Sigma, 5438040100) was added for a final concentration of 2% (v/v). Samples were centrifuged at 11,000 g for 5 min. 5% of the supernatant was used for stage tipping. C_18 S_tageTips were made as described in Rappsilber *et al.*^79^. StageTips were washed with sequential washes of 50 μL HPLC-grade methanol (Sigma, 494291), Buffer B, and Buffer A, centrifuging the tips at 1,800 g for 2 min between each wash. Buffer B consisted of 0.1% HPLC-grade trifluoroacetic acid (TFA, Alfa Aesar, AA446305Y) and 80% HPLC-grade acetonitrile (Sigma-Aldrich, 271004). Buffer A consisted of 0.1% TFA and 5% acetonitrile. Samples were loaded on the StageTips, centrifuged at 1,800 g for 2 min, and washed again with Buffer A.

### LC-MS/MS-based proteomics and data analysis

Peptide samples were separated on an EASY-nLC 1200 System (Thermo Fisher Scientific) using 20 cm long fused silica capillary columns (100 µm ID, laser pulled in-house with Sutter P-2000, Novato CA) packed with 3 μm 120 Å reversed phase C18 beads (Dr. Maisch, Ammerbuch, DE). The LC gradient was 90 min long with 5−35% B at 300 nL/min. LC solvent A was 0.1% (v/v) aq. acetic acid and LC solvent B was 20% 0.1% (v/v) acetic acid, 80% acetonitrile. MS data was collected with a Thermo Fisher Scientific Orbitrap Fusion Lumos. The MS method used was a data-independent acquisition (DIA) method with a 120K resolution MS1 scan from m/z 395-1005 and 15K resolution MS2 scans with 12 Th isolation with between m/z 400-1000; the AGC target was 400,000 ions with a max injection time of 25 ms. Raw files were analyzed with MSFragger 3.4^80^ and DIA-nn 1.8^81^ using protein and peptide FDRs of 0.01. The FASTA database used for search was a human Uniprot reference database downloaded 2021-11-08, with ‘stricttrypsin’ and 2 missed cleavages permitted. Variable modifications were N-term acetylation, pyroGlu- and loss of ammonia (-17.0265 nQnC) at peptide N-termini. Carbamidomethyl (C) was a fixed modification. Data was further processed using the Perseus software package (version 1.5.2.6), the R environment, Origin Pro 8.0 and Microsoft Excel. GO term enrichment analysis was performed using the ShinyGO 0.76 web application (http://bioinformatics.sdstate.edu/go/)^36^. LC-MS/MS data accession numbers are: MassIVE MSV000096493, ProteomeXchange PXD058152.

### RNA sequencing

500,000 HeLa cells were plated in 6-well plates. After two days, total RNA was isolated using an RNeasy kit (Qiagen, 74106). Samples were sent to Genewiz for sequencing and data analysis, as follows: Samples underwent mRNA enrichment, mRNA fragmentation, and random priming. Genewiz performed first and second strand cDNA synthesis followed by end repair, 5’ phosphorylation, and dA-Tailing. cDNA was sequenced, following adaptor ligation and enrichment by PCR. During analysis, the reads were trimmed and mapped to the Homo sapiens GRCh38 reference genome available^82^ (https://www.ensembl.org) using the STAR aligner (v.2.5.2b). Hit counts for genes and exons were calculated with featureCounts (Subread package, v.1.5.2). DESeq2 was used for analysis of differential gene expression. p-values and log_2 f_old changes were determined with the Wald test, and genes with an adjusted p-value < 0.05 and absolute log_2 f_old change > 1 were considered to be differentially expressed.

### Seahorse metabolic flux analysis

HeLa and AML12 cells were seeded at 10,000 or 15,000 cells/well, respectively, in a XFe96 Microplate (Agilent Technologies, 103792-100) and cultured overnight at 37°C in 5% CO_2_. The Extracellular Flux Assay Kit oxygen probes (Agilent Technologies, 103792-100) were activated in XF calibrant buffer (Agilent, 100840-000) at 37°C in a non-CO_2 i_ncubator overnight. Seahorse XF medium was prepared fresh by supplementing XF DMEM Medium, pH 7.4 (Agilent, 103575-100) with 100 mM Pyruvate Solution (Agilent, 103578-100), 1 M Glucose Solution (Agilent, 103577-100), and 200 mM L-Glutamine (Fisher Scientific, BP379-100). The cell culture medium was removed from the plate and the cells were washed once with pre-warmed Seahorse XF medium and replaced with fresh medium a second time. The cells were then incubated in a CO_2-_free incubator at 37°C for 1 hr. Oxygen consumption rate (OCR) measurements were performed with a Seahorse XFe96 Extracellular Flux Analyzer (Agilent, S7800B) using the Cell Mito Stress Test program. The sensor cartridge was loaded with drugs and inserted into the machine 30 min prior to the flux assay to calibrate the oxygen probes. Following measurements of baseline cellular respiration, respiration was measured after addition of oligomycin, carbonyl cyanide-p-trifluoromethoxyphenylhydrazone (FCCP), and antimycin A and rotenone together, with final concentrations of 1 μM. Oxygen consumption rate was analyzed with the Seahorse XF Cell Mito Stress Test Report Generator (Wave Software). OCR measurements were normalized to cell number as determined by the CyQUANT Cell Proliferation Assay Kit (Thermo Scientific, C7026) or DAPI staining for HeLa and AML12 cells, respectively. For the CyQUANT assay, the media was removed from each well and the plates were frozen at -80°C for >1 hr to lyse the cells. CyQUANT was utilized per manufacturer’s instructions to measure fluorescence intensity (Ex 480 nm, Em 520 nm). For DAPI staining, 40 μL media was left in each well and cells were frozen at -80°C for >1 hr to lyse the cells. After thawing, 100 μL 1 μg/μL DAPI (Thermo Scientific, 62248) was added to each well for 5 min before measuring DAPI fluorescence (Ex 359 nm, Em 457 nm).

#### LC-MS-based lipid analysis

Cell pellets containing 10 million HeLa cells per condition were frozen at -80°C until lipid extraction. Lipid extraction and analysis were carried out as described^83^. Briefly, cell pellets were resuspended in methanol and sonicated 3×30 sec while on ice. The homogenized solution was centrifuged at 16,900 g for 15 min at 4°C. 800 μL of supernatant was removed into a fresh 4-dram vial. 1 mL cold methanol was added to the remaining pellet, sonicated, and centrifuged again. This supernatant was combined with the previous extract and dried down using a nitrogen evaporator. LC-MS data was acquired using an Agilent 1260 HPLC with an Agilent 6530 Quadrupole Time-of-Flight mass spectrometer. A Luna C5 reverse-phase column (5m, 4.6 x 50mm, Phenomenex) was used for positive mode and a Gemini C18 reversed-phase column (5m, 4.6 x 50mm, Phenomenex) was used in negative mode. Mobile phase A was 95:5 water:methanol and mobile phase B was 60:35:5 isopropanol:methanol:water for both negative and positive modes. The mobile phases contained additives to aid electrospray ionization and improve ion detection (0.1% formic acid in positive mode; 5 mM ammonium formate, and 0.1% ammonium hydroxide in negative mode). Data acquisition in positive and negative mode used different gradients: ramping from 0% mobile phase B to 100% mobile phase B occurred over 45 min and 65 min for positive and negative mode, respectively. For both methods, the flow rate was set to 0.1 mL/min for the first 5 min at 100% mobile phase A then increased to 0.5 mL/min for the remainder of the run. A dual electrospray ionization source was used; capillary and fragmentor voltages were set to 3500 V and 175 V, respectively. All data were collected in the extended dynamic range mode (*m/z* = 50 - 1,700). Representative lipid species for each class were targeted by extracting the corresponding m/z for each ion in MassHunter Qualitative Analysis software (version B.06.00, Agilent Technologies). Peak areas were manually integrated and represented as abundance. Relative abundance values were calculated by dividing the abundance of a lipid by the average abundance of that lipid in the wildtype set.

### Cell lysate preparation and immunoblots

Cells were grown to 40-80% confluency, then plates were washed with cold PBS (Thermo Fisher Scientific, 20012050) on ice and lysed with either RIPA or triton lysis buffer. RIPA buffer consisted of 50 mM HEPES-KOH, pH 7.4 (Sigma-Aldrich, H3375; Sigma Millipore, 1050121000), 150 mM NaCl (Sigma-Aldrich, 746398), 2 mM EDTA (Sigma-Aldrich, 607-429-00-8), 10 mM sodium pyrophosphate (Sigma-Aldrich, 71501), 10 mM β-glycerophosphate (Sigma-Aldrich, G9422), 1% sodium deoxycholate (Sigma-Aldrich, D6750), 0.1% SDS (Sigma-Aldrich, L4509), and 1% Triton X-100 (Sigma-Aldrich, X100). Triton buffer consisted of 50 mM HEPES-KOH, pH 7.4, 40 mM NaCl, 2 mM EDTA, 10 mM β-Glycerophosphate, 1.5 mM sodium orthovanadate (Sigma-Aldrich, S6508), 50 mM sodium fluoride (Sigma-Aldrich, S7920), 10 mM sodium pyrophosphate, and 1% Triton X-100. Lysis buffers were supplemented with protease inhibitors (Sigma-Aldrich, 11836170001) immediately prior to lysis. Lysates were centrifuged at 17,000 g for 10 min at 4°C. The supernatants were normalized by protein concentration as measured by either a Bradford protein assay (Bio-Rad, 5000205) or Pierce BCA assay (Thermo Fisher Scientific, 23209) and a BioTek Synergy H1 plate reader (Fisher Scientific, 11-120-536). SDS-PAGE samples were prepared from the lysates with 5X reducing sample buffer (pH 6.8) consisting of 10% SDS, 25% 2-mercaptoethanol (Sigma-Aldrich, M3148), 25% glycerol (Sigma-Aldrich, G5516), 50 mM Tris-HCl (Sigma-Aldrich, RDD008), and 0.1% bromophenol blue (VWR, 97061-690).

Samples were heated to 95°C for 5 min and then brought to room temperature. Samples were mixed by pipetting and loaded onto 4-15% gradient Mini-PROTEAN TGX gels (Bio-Rad, 4561086). Following gel electrophoresis in Tris-Glycine-SDS running buffer (Boston BioProducts, BP-150), proteins were transferred to ethanol-activated 0.2 μm PVDF membranes (Bio-Rad, 1620174) using the BioRad Mixed Molecular Weight Protein Transfer setting (7 min, 1.3 A, 25 V) on the Trans-Blot Turbo Transfer System (Bio-Rad, 1704150) with Trans-Blot Turbo Transfer Buffer (Bio-Rad, 10026938). After transfer, membranes were washed in 20% ethanol for ∼2 min and checked for equal loading and even transfer by Ponceau stain (Sigma-Aldrich, P7170). Membranes were then incubated with 1% wt/vol BSA (Sigma-Aldrich, A2153) in TBST with 1% (v/v) Tween-20 (Boston Bio-Products, IBB-180) at room temperature for 1-4 hr. Membranes were incubated overnight with primary antibody and 1% BSA in TBST at 4°C. All membranes were then washed with TBST three times, 5 min each, and incubated with secondary antibody and 1% BSA in TBST for 1 hr using a BlotCycler W5 (Precision Biosystems). Membranes were washed 4 times, 5 min each, with TBST. Membranes were developed using Clarity Western ECL substrate (Bio-Rad, 170-5060) or Clarity Max Western ECL substrate (Bio-Rad, 1705062). Immunoblots were imaged with an iBrightCL1000 imager (Life Technologies, A23749) and quantified using the Image Studio Lite software, version 5.2 (LI-COR Biosciences). The following antibodies and dilutions were used: 1:3000 rabbit ACAT1 antibody (ABclonal, A13273); 1:3000 rabbit BCKDHA antibody (ABclonal, A21588); 1:5000 mouse β-actin antibody (Cell Signaling Technology, 3700); 1:5000 rabbit EMRE antibody (Bethyl Laboratories, A300-BL19208); 1:3000 rabbit HSP60 antibody (Cell Signaling Technology, 12165); 1:3000 rabbit KLF15 antibody (ABclonal, A7194); 1:3000 rabbit MCEE antibody (ABclonal, A14430); 1:3000 rabbit MCU antibody (Cell Signaling Technology, 14997); 1:3000 rabbit MICU1 antibody (ABclonal, A21948); 1:3000 rabbit MMSDH antibody (ABclonal, A3309); 1:3000 rabbit MMUT antibody (ABclonal, A3969); 1:1000 mouse OTC antibody (Invitrogen, PAS-28197); 1:3000 rabbit PCCB antibody (ABclonal, A5415); 1:3000 rabbit phospho-BCKDH-E1α, human S292/mouse S293 antibody (Cell Signaling Technology, 40368); 1:1000 rabbit phospho-R-X-S*/T* (Cell Signaling Technology, 9621); 1:3000 rabbit PKA-Cα antibody (BD Biosciences, 610980); 1:3000 rabbit TOM20 antibody (Cell Signaling Technology, 42406); 1:10,000 HRP-linked α-rabbit secondary antibody (Cell Signaling Technology, 7074); 1:10,000 HRP-linked α-mouse secondary antibody (Cell Signaling Technology, 7076).

### Isolation of patient samples

Fresh non-tumor liver (NTL) and FLC tumor samples were procured from patients undergoing liver resection for FLC treatment. Prior to surgery, written-informed consent was obtained for tissue donation under a research protocol approved by the University of Washington Institutional Review Board (IRB) (#1852) and Seattle Children’s Hospital IRB (#15277). All research samples were de-identified. Clinical details for each patient are outlined in Supplemental Table 1. NTL and FLC samples were processed and transferred to the laboratory on ice within 1 hr of tissue excision. Samples were stored at -80°C prior to lysate preparation.

### Lysis and sample preparation of patient samples

Tissues were lysed with protease inhibitor-supplemented RIPA buffer using a TissueRuptor homogenizer (Qiagen, 9002755). Tissues were homogenized until they were a uniform consistency. Samples were then centrifuged at 17,000 g for 10 min at 4°C and prepared for SDS-PAGE with 5X reducing sample buffer.

### Generation of shRNA constructs

shRNA sequences were designed by the RNAi Consortium (TRC) of the Broad Institute. Forward and reverse oligos encoding the shRNA were annealed to form dsDNA. 100 pmol forward oligo and 100 pmol reverse oligo were resuspended in 25 μL 1X NEBuffer 2 (NEB, B7002S). The solution was incubated in a 95°C water bath for 4 min then slowly cooled (∼0.5°C/min) to room temperature. The pLKO.1 vector (Addgene, 10878) was digested with AgeI-HF (NEB, R3552) and EcoRI-HF (NEB, R3101) and ligated with the oligo solution according to the NEB Quick Ligase (NEB, M2200) protocol. The ligation mixture was transformed into XL10-Gold competent cells (Agilent, 200315). Resulting plasmids were verified by sequencing.

Mouse shMCU-1, forward oligo: CCGGGATCCGAGATGACCGTGAATCCTCGAGGATTCACGGTCATCTCGGATCTTTTTG;

reverse oligo: AATTCAAAAAGATCCGAGATGACCGTGAATCCTCGAGGATTCACGGTCATCTCGGATC.

Mouse shMCU-2, forward oligo: CCGGTAGGGAATAAAGGGATCTTAACTCGAGTTAAGATCCCTTTATTCCCTATTTTTG;

reverse oligo: AATTCAAAAATAGGGAATAAAGGGATCTTAACTCGAGTTAAGATCCCTTTATTCCCTA. shGFP

control, forward oligo: CCGGCCACATGAAGCAGCACGACTTCTCGAGAAGTCGTGCTGCTTCATGTGGTTTTTG;

reverse oligo: AATTCAAAAACCACATGAAGCAGCACGACTTCTCGAGAAGTCGTGCTGCTTCATGTGG

shRFP control, forward oligo: CCGGGCTCCGTGAACGGCCACGAGTCTCGAGACTCGTGGCCGTTCACGGAGCTTTTTG;

reverse oligo: AATTCAAAAAGCTCCGTGAACGGCCACGAGTCTCGAGACTCGTGGCCGTTCACGGAGC

### Lentivirus production and transduction

1 million HEK293T cells (from the Whitehead Institute for Biomedical Research) were plated in 6 cm plates. The following day, cells were transfected. 100 ng p-CMV-VSV-G (Addgene, 8454), 900 ng psPax2 (Addgene, 12260), and 1 μg viral plasmid were diluted in 6 μL X-tremeGENE 9 DNA transfection reagent (Sigma-Aldrich, 6365787001) and 150 μL DMEM and incubated at room temperature for 15 min before being added to the cells. After 36-48 hr, the virus-containing medium was filtered through a 0.45 μm sterile filter (VWR, 28145-505) and stored at -20°C until use. 200,000 HeLa or 100,000 AML12 cells were plated in 6-well plates. The following day, 200 μL of the virus-containing media and 8 μg/mL polybrene (Sigma-Aldrich, H9268) was added to the cells. Infected cells were selected for 48 hr with 2 μg/mL puromycin. For assays measuring transcriptional changes in the BCAA catabolism pathway, shMCU-infected cells were cultured for 2-3 weeks prior to harvesting.

### RNA isolation, cDNA synthesis, qPCR

AML12 cells were grown in 6-well plates to ∼80% confluency in triplicate. Tumor and non-tumor liver samples were stored in RNALater solution at -80°C prior to RNA extraction. RNA was isolated with the RNeasy Kit (Qiagen, 74106). For liver samples, ∼30 mg of tissue was homogenized using a TissueRuptor homogenizer (Qiagen, 9002755) according to the kit protocol. 100 ng of RNA was used for cDNA synthesis using the Maxima First Strand cDNA synthesis kit (Thermo Fisher Scientific, K1641) according to kit instructions. The resulting cDNA was diluted 1:30 in dH_2O_, and either 1 or 2 μL of diluted cDNA was used for qPCR using Applied Biosystems TaqMan Fast Advanced Master Mix (Thermo Fisher Scientific, 44-445-57) in a 384-well plate (Thermo Fisher Scientific, 43-098-49) using a QuantStudio 5 RT-PCR System (Thermo Fisher Scientific). The following Taqman probes were used: human ACTB (Thermo Fisher Scientific, 4331182-Hs01060665_g1), human MCU (Thermo Fisher Scientific, 4331182-Hs00293548_m1), and human OTC (Thermo Fisher Scientific, 4331182-Hs00166892_m1).

### Immunohistochemistry

Fresh human FLC tissue from liver resections were fixed in formalin and paraffin-embedded. 4 µm thick sections were cut and placed on glass slides. Slides were stained with Hematoxylin and Eosin (H&E) using a H&E staining kit (Vector Laboratories, H-3502) according to supplied protocol. For immunohistochemistry staining, slides were deparaffinized, rehydrated, and washed with TBST. After antigen retrieval in 10 mM sodium citrate (pH 6) and quenching of endogenous peroxidase activity with 3% H_2O2_, samples were blocked with 2.5% normal horse serum (NHS) before incubation with primary antibodies overnight at 4°C. The following primary antibodies were used: 1:300 KLF15 rabbit antibody (ABclonal, A7194) and 1:250 MCU rabbit antibody (Cell Signaling Technologies, 14997). Negative controls were treated with 2.5% NHS without primary antibodies. Signals were processed using the ImmPRESS HRP Horse Anti-Mouse (Vector Laboratories, MP-7402) and Anti-Rabbit (Vector Laboratories, MP-7401) Peroxidase Kits and ImmPACT DAB Substrate Kit Peroxidase (Vector Laboratories, SK-4105) according to manufacturer’s protocol. Slides were counterstained with Hematoxylin QS, dehydrated, and mounted with VectaMount Express Mounting Medium.

### Ca^2+^ uptake in permeabilized cells

Protocol is adapted from Sancak *et al.*^10^. Cell medium was changed 1-2 hr prior to harvesting. Cells were detached with trypsin (Gibco, 12605-010) and resuspended in cell medium. 1 million HeLa cells or 0.5 million AML12 cells were centrifuged at 800 g for 3 min. Cell pellets were washed with PBS (Thermo Fisher Scientific, 20012050), centrifuged at 800 g for 3 min, and resuspended in 150 μL KCl buffer. KCl buffer consists of 125 mM KCl (Sigma-Aldrich, 793590), 2 mM K_2H_PO_4 (_Sigma-Aldrich, P3786), 1 mM MgCl_2_, (Sigma-Aldrich, M8266), and 20 mM HEPES, pH 7.2 (Sigma-Aldrich, H3375) supplemented fresh with 5 mM glutamate (Sigma-Aldrich, G1251), 5 mM malate (Sigma-Aldrich, M7397), and 1 μM Oregon Green 488 Bapta-6F (Invitrogen, O23990). 0.01% or 0.005% digitonin (Thermo Fisher Scientific, BN2006) was added for HeLa and AML12 cell assays, respectively. Cells were transferred to a black 96-well plate (Greiner Bio-One, 655076). Fluorescence (Ex 485/20 nm, Em 520/20 nm) was monitored every 2 sec for 136 sec at room temperature using a BioTek Synergy H1 microplate reader (Fisher Scientific, 11-120-536) before and after injection of 50 μM CaCl_2 (_Sigma-Aldrich, 746495). Ca^2+^ uptake rates were calculated using the linear fit of uptake data points between 20 and 30 sec after Ca^2+^ injection. A maximum of eight samples were assayed together, including one wildtype control per assay run.

### Ca^2+^ release from permeabilized cells

Cell medium was changed 1-2 hr prior to harvesting. Cells were detached with trypsin (Gibco, 12605-010) and resuspended in cell medium. 1 million HeLa cells or 0.5 million AML12 cells were centrifuged at 800 g for 3 min. Cell pellets were washed with PBS (Thermo Fisher Scientific, 20012050), centrifuged at 800 g for 3 min, and resuspended in 150 μL KCl buffer. KCl buffer consists of 125 mM KCl (Sigma-Aldrich, 793590), 2 mM K_2H_PO_4 (_Sigma-Aldrich, P3786), 1 mM MgCl_2_, (Sigma-Aldrich, M8266), and 20 mM HEPES, pH 7.2 (Sigma-Aldrich, H3375) supplemented fresh with 1 μM Oregon Green 488 Bapta-6F (Invitrogen, O23990). 0.01% or 0.005% digitonin (Thermo Fisher Scientific, BN2006) was added for HeLa and AML12 cell assays, respectively. Cells were transferred to a black 96-well plate (Greiner Bio-One, 655076). Fluorescence (Ex 485/20 nm, Em 520/20 nm) was monitored every 2 sec for 196 sec at room temperature using a BioTek Synergy H1 microplate reader (Fisher Scientific, 11-120-536) before and after injection of 1 μM CCCP (Cayman Chemical Company, 25458) in KCl buffer at 12 sec. Ca^2+^ release was calculated using the absolute difference between fluorescence reads prior to CCCP injection (average of six values, 0-10 sec) and 3 min after injection (average of six values, 186-196 sec). A maximum of eight samples were assayed together, including one wildtype control per assay run. For PKA kinase inhibition assays, AML12 cells were cultured for 4 days with 5 μM BLU2864 or DMSO vehicle control. Cells were passaged on day 2, and fresh medium and drug were added. Ca^2+^ release experiments with BLU2864 were performed with ∼0.5 million cells, however BLU2864-treated cells formed clumps and could not be counted precisely. Data were instead normalized by protein concentration. Following Ca^2+^ release, 1 μL Triton X-100 (Sigma-Aldrich, X100) was added to each well, and the solution was pipetted vigorously. Protein concentration was then determined by Bradford assay (Bio-Rad, 5000205).

### TEM of FLC patient liver

Tissue fixation was performed with 4% glutaraldehyde in 0.1 M sodium cacodylate buffer, then stored overnight at 4 °C. The tissue was then washed 5 x 5 min in buffer at room temperature and post fixed in buffered 2% osmium tetroxide on ice for 1 hr. This was followed by 5 washes in ddH_2O_, then en bloc staining in 1% uranyl acetate (aqueous), overnight at 4 °C. The following day the tissue was washed 5 x 5 minutes in ddH_2O_ then dehydrated in ice cold 30%, 50%, 70%, and 95% ethanol, then allowed to come to room temperature. This was followed by 2 changes of 100% ethanol and two changes of propylene oxide. The tissue was then infiltrated in a 1:1 mixture of propylene oxide:Epon Araldite resin for 2 hr followed by two changes of fresh Epon Araldite, 2 hr each change. It was then placed in flat embedding molds and polymerized at 60 °C overnight. Fixed samples were sliced to a thickness of 80 nm and were imaged on a JEOL1230 Transmission Electron Microscope (TEM) operated at 80 kV.

### Quantification of Mitochondria and deposits from EM images

Images of patient liver tissue taken at 10,000x magnification were analyzed by two independent, blinded researchers. First, mitochondria were identified in each image. Mitochondria were characterized as structures with a well-defined, curved border, typically 0.5-1 µm in diameter, with a mottled, gray matrix and at least one visible cristae line. Confirmed mitochondria were examined for the presence or absence of black, electron-dense deposits. Deposits were defined as speckles that were both darker and thicker in diameter than cristae membrane lines. With FLC patient 42.2, for which no non-tumor liver sample was available, cells from the tumor periphery were classified as oncocytic or peri-oncocytic. Cells with a circular nucleus and dispersed mitochondria were classified as peri-oncocytic; cells with misshapen nuclei and abundant cytoplasmic mitochondria were classified as oncocytic.

### TMRM measurements

500,000 AML12 cells were centrifuged at 800 g for 3 min, washed with 1 mL PBS (Thermo Fisher Scientific, 20012050), and centrifuged again at 800 g for 3 min. The supernatant was aspirated and the cell pellet was resuspended in 150 μL KCl buffer (for composition, see Ca^2+^ uptake protocol above) supplemented with 500 nM tetramethylrhodamine methyl ester (TMRM, Thermo Fisher Scientific, I34361) and 0.005% digitonin (Thermo Fisher Scientific, BN2006). The cell suspension was transferred to a black 96-well plate (Greiner Bio-One, 655076). Fluorescence measurements (Ex 540 nm, Em 590 nm) were taken across three periods using a BioTek Synergy H1 microplate reader (Fisher Scientific, 11-120-536). The first measurements were taken every second for 1 min to establish a baseline. Then 500 mM glutamate (Sigma-Aldrich, G1251) and 5 mM malate (Sigma-Aldrich, M7397) stocks were injected for final concentrations of 5 mM, and a second reading was taken every second for 1 min to obtain the minimum fluorescence, corresponding with maximal mitochondrial membrane potential. Finally, mitochondrial membrane potential was dissipated with the addition of 1 μM CCCP (Cayman Chemical Company, 25458), and reads were taken every second for 2 min to obtain the maximum fluorescence. Relative mitochondrial membrane potential was reported as the difference between the maximum (average of data from 294-304 sec) and minimum TMRM fluorescence measurements (average of data from 172-182 sec).

### Generation of mito-GECO constructs and matrix and cytosolic Ca^2+^ measurements

The pLMM1 vector was generated from pLYS1^84^ by replacing the CMV promoter with the EF1α promoter. G-GECO1.1^46^ was cloned into either pLYS1 or pLMM1 with an N-terminal mitochondrial targeting sequence taken from cytochrome C subunit 8A (COX8). AML12 cells were transduced with mito-G-GECO lentiviral constructs generated using pLYS1 (WT and c14) or pLMM1 (c4) and selected with puromycin as described above. 80,000 AML12 wildtype, c14, and c4 cells expressing mito-G-GECO were cultured in 12-well plates for 2 days until reaching 90-95% confluency. Medium was then aspirated and replaced with 500 μL phenol red-free DMEM/F-12 (Thermo Scientific, 21041025) supplemented with 10% FBS (VWR, 89510-186) and 100 IU/mL penicillin and 100 μg/mL streptomycin (VWR, 45000-652) for 1.5 hr. Fluorescence measurements (Ex 480 nm, Em 520 nm) were obtained with a BioTek Synergy H1 microplate reader (Fisher Scientific, 11-120-536) at 37°C. To establish baseline, fluorescence was measured every 10 sec for 2 min. Then CCCP was added to a final concentration of 50 μM (Cayman Chemical Company, 25458) to each well, and fluorescence was measured to determine minimum fluorescence. Finally, 5 μL of 1mM stock of ionomycin (Sigma-Aldrich, I3909) and 10 μL of 1M stock of CaCl_2 w_ere added, and fluorescence was measured to establish maximum fluorescence. To correct for potential differences in mito-G-GECO expression across cell lines, baseline results were reported as a percentage of the difference between maximum and minimum values. To calculate the baseline fraction, minimum fluorescence is subtracted from baseline fluorescence over minimum fluorescence subtracted from maximum fluorescence. For each measurement, baseline, minimum, and maximum values were determined manually.

AML12 cells were transduced with cyto-G-GECO lentiviral constructs generated using pLYS1 and selected with puromycin as described above. 12-well plates were coated with EmbryoMax® 0.1% Gelatin Solution (Sigma-Aldrich, ES-006) for 10 min. Excess gelatin solution was aspirated off, and 80,000 AML12 wildtype, c14, and c4 cells expressing cyto-G-GECO were cultured in the 12-well plates for 2 days until reaching 90-95% confluency. Medium was then aspirated and replaced with 500 μL phenol red-free DMEM/F-12 (Thermo Scientific, 21041025) supplemented with 10% FBS (VWR, 89510-186) and 100 IU/mL penicillin and 100 μg/mL streptomycin (VWR, 45000-652) for 1.5 hr. Fluorescence measurements (Ex 480 nm, Em 520 nm) were obtained with a BioTek Synergy H1 microplate reader (Fisher Scientific, 11-120-536) at 37°C. To establish baseline, fluorescence was measured every 10 sec for 2 min. Then 5 μM ionomycin (Sigma-Aldrich, I3909) and 2.5 μM EGTA (Sigma-Aldrich, E3889) were added to each well, and fluorescence was measured to determine minimum fluorescence. Finally, 10 μM CaCl_2 w_as added, and fluorescence was measured to establish maximum fluorescence. To correct for potential differences in cyto-G-GECO expression across cell lines, baseline results were reported as a percentage of the difference between maximum and minimum values. To calculate the baseline fraction, minimum fluorescence is subtracted from baseline fluorescence over minimum fluorescence subtracted from maximum fluorescence. For each measurement, baseline, minimum, and maximum values were determined manually.

### ^13^C_6_ Leucine Carbon Tracing

800,000 Hela WT and MCU KO cells were plated in two 6-well plates to achieve 85-95% confluency. The next day, one plate was taken and washed twice with PBS (Thermo Fisher Scientific, 20012050), and the media was changed to DMEM/F-12 without amino acids (US Biological, D9807-02A, prepared with 84 g/mol of sodium bicarbonate, 238 g/mol of HEPES, and 180 g/mol of glucose, pH to 7.2) mixed with 10mM L-Leucine-^13^C_6 (_MedChem Express, HY-N0486S2). This DMEM/F-12 w/o glucose or amino acids was supplemented with dialyzed 10% FBS (VWR, 89510-186), 1X insulin-transferrin-selenium (Gibco, 41400045), 40 ng/mL dexamethasone (MP Biomedicals, 0219456180), and 100 IU/mL penicillin and 100 μg/mL streptomycin (VWR, 45000-652) and all amino acid (except leucine) concentrations matching that of DMEM/F-12 (Thermo Scientific, 21041025). FBS dialysis was done with Slide-A-Lyzer™ Dialysis Flasks (ThermoFisher Scientific, 87761) to 3.5kD. This plate was then left to incubate in 37°C with 5% CO_2 f_or 2 hrs to ensure that L-Leucine-13C6 was incorporated. The plate was then quickly washed with 1X HPLC-grade ice cold saline (Fisher, 23293184) which was aspirated before adding 500 μL ice cold 80% HPLC-grade methanol (Sigma, 646377) in HPLC-grade water (Sigma, 270733). Each well was quickly scraped using a cell scraper to disrupt the cell layer, quenching metabolism and preventing significant amounts of evaporation. Depending on evaporation, 400-500 μL of liquid in each well was transferred to a microcentrifuge tube, and this was centrifuged at 17,000 g for 10 min at 4°C. As much of the same amount of supernatant was then taken from each and transferred to a new microcentrifuge tube and this was centrifuged again at 17,000 g for 10 min at 4°C. Finally, 350 μL of the supernatant of each sample was transferred to a new microcentrifuge tube and lyophilized. After collection, the second 6-well plate was taken and counted using a Coulter Z2 Cell and Particle Counter (Beckman Coulter, 383550) to obtain cell counts for normalization of each sample.

### Liquid Chromatography-Mass Spectrometry (LC-MS)

Lyophilized samples were resuspended in 80% HPLC grade methanol in HPLC grade water and transferred to liquid chromatography-mass spectrometry (LCMS) vials for measurement by LCMS. Metabolite quantitation was performed using a Q Exactive HF-X Hybrid Quadrupole-Orbitrap Mass Spectrometer equipped with an Ion Max API source and H-ESI II probe, coupled to a Vanquish Flex Binary UHPLC system (Thermo Scientific). Mass calibrations were completed at a minimum of every 5 days in both the positive and negative polarity modes using LTQ Velos ESI Calibration Solution (Pierce). Polar Samples were chromatographically separated by injecting a sample volume of 1 μL into a SeQuant ZIC-pHILIC Polymeric column (2.1 x 150 mm 5 mM, EMD Millipore). The flow rate was set to 150 mL/min, autosampler temperature set to 10 °C, and column temperature set to 30 °C. Mobile Phase A consisted of 20 mM ammonium carbonate and 0.1% (v/v) ammonium hydroxide, and Mobile Phase B consisted of 100 % acetonitrile. The sample was gradient eluted (% B) from the column as follows: 0-20 min.: linear gradient from 85% to 20% B; 20-24 min.: hold at 20% B; 24-24.5 min.: linear gradient from 20% to 85% B; 24.5 min.-end: hold at 85% B until equilibrated with ten column volumes. Mobile Phase was directed into the ion source with the following parameters: sheath gas = 45, auxiliary gas = 15, sweep gas = 2, spray voltage = 2.9 kV in the negative mode or 3.5 kV in the positive mode, capillary temperature = 300°C, RF level = 40%, auxiliary gas heater temperature = 325°C. Mass detection was conducted with a resolution of 240,000 in full scan mode, with an AGC target of 3,000,000 and maximum injection time of 250 msec. Metabolites were detected over a mass range of 70-1050 m/z. Quantitation of all metabolites was performed using Tracefinder 4.1 (Thermo Scientific) referencing an in-house metabolite standards library using ≤ 5 ppm mass error. Data from leucine isotope labeling experiments includes correction for natural isotope abundance using IsoCor software v.2.2.

### NADH/NAD^+^ measurements

20,000 Hela WT and MCU KO cells were plated in 6-well plates with DMEM media (Thermo Fisher Scientific, 11-965-118) supplemented with 10% FBS (VWR, 89510-186), 2 mM GlutaMAX (Thermo Fisher Scientific, 35-050-061), 100 IU/mL penicillin, and 100 μg/mL streptomycin (VWR, 45000-652) and allowed to adhere overnight. The medium was aspirated and rinsed with PBS (Thermo Fisher Scientific, 20012050), and replaced with DMEM:F12 w/o Glucose, Amino Acids (US Biological, D9807-02A) that was supplemented with dialyzed 10% FBS, 100 IU/mL penicillin and 100 μg/mL streptomycin, glucose, and either non-branched-chain amino acids or all amino acids. Plates were incubated for 3 hr at 37°C in 5% CO_2_. Cells were washed three times in ice-cold PBS. They were then extracted in 100 μL of ice-cold lysis buffer (1% dodecyltrimethylammonium bromide [DTAB] in 0.2 N of NaOH diluted 1:1 with PBS), and immediately frozen at −80°C and kept there until all replicates were collected. For NADH measurement, 40 μL of freshly thawed lysates was transferred to microcentrifuge tubes and incubated at 75°C for 30 min for base-mediated degradation of NAD^+^. For NAD^+^ measurement, 40μL of lysis buffer was added with 40 μL of 0.4N HCl and 40 μL of freshly thawed lysate in microcentrifuge tube. This was then incubated at 60°C for 15 min for acidic conditions to selectively degrade NADH. After respective incubations, samples were allowed to equilibrate to room temperature for 8 minutes. NADH measurement samples were quenched with 40μL of 0.25 M Tris in 0.2 N HCl. NAD^+^ measurement samples were quenched with 40 μL of 0.5 M Tris base. 0 nM, 10 nM, 100 nM, 200 nM, 400 nM stocks were prepared right before running the assay by diluting 1 μM NAD^+^ in a mixture of equal volumes of PBS, base solution with 1% DTAB, 0.4 N HCl, and 0.5 M Trizma base. The working reagent (reconstituted luciferan detection reagent, reductase, reductase substrate, NAD^+^ cycling enzyme, and NAD cycling substrate) added to the NADH and NAD^+^ samples were prepared using the instructions provided by the NAD^+^/NADH Glo Assay Kit (Promega, G9071). 1:1 of sample or standard solution and working reagent from kit were added to a plate using multichannel pipette. Enzyme-linked luminescence-based NAD+ and NADH measurement were measured at 0 and 40 minutes using the BioTek Synergy H1 plate reader (Fisher Scientific, 11-120-536). Luminescence readings of NAD^+^ and NADH was used to calculate the NADH/NAD^+^ ratio and subsequent fold changes.

### Analysis of Gene Expression

Raw data files (FASTQ format) were downloaded from the European Nucleotide Archive for each GEO study. Counts per gene were computed from the FASTQ files by first aligning to the NCBI GRCh38 transcriptome using the salmon aligner^85^, and then collapsing to the gene level using the Bioconductor tximport package^86^. The Bioconductor edgeR^87^ package was then used to fit linear models for each study using the limma-voom pipeline^86^, and then computing empirical Bayes adjusted contrasts^88^ between FLC and NML. For those datasets with complete pairing, the linear model included a blocking effect for patient, and for those with only partial pairing a linear mixed effects model was fit to account for within-patient correlation. A meta-analysis was then performed using Stouffer’s method^89^, implemented in the Bioconductor metapod package^90^, using the inverse of the effect size as weights to improve precision^91^.

## ACKNOWLEDGEMENTS

We are thankful to the patients who donated samples to be used in research. We are also grateful to Dr. Vishal Gohil and for his thoughtful discussion of this project. We thank Edward Parker for his help with processing patient samples for electron microscopy and image acquisition; Dr. Rong Tien for sharing KLF15 antibody and plasmids; Dr. Taran Gujral and his lab for useful discussion, guidance and help with initial data analysis. This work used an EASY-nLC1200 UHPLC and Thermo Scientific Orbitrap Fusion Lumos Tribrid mass spectrometer purchased with funding from a National Institutes of Health SIG grant S10OD021502. Several figures were created with tools from BioRender.com. This research was supported by the Proteomics & Metabolomics Shared Resource, which is supported by a National Cancer Institute Cancer Center Support Grant for the Fred Hutch/University of Washington/Seattle Children’s Cancer Consortium (NCI; P30CA015704), R35 GM136234 (Y.S.), Pew Charitable Trusts (Y.S.), 1F31AG072716-01A1 (M.J.S.M), NSF MCB2314338 (G.E.A.G), P30 CA015704 (R.S.Y, H.L.K), DOD W81XWH1910544 (R.S.Y), R01GM129090 (S.E.O), R35GM147118 (L.B.S.), RO1CA279997 (J.D.S), Fibrolamellar Foundation (J.D.S) The University of Washington EDGE center is supported by NIH P30ES007033.

## AUTHOR CONTRIBUTIONS

- NM Marsh: conceptualization, data collection, data analysis, validation, visualization, methodology, and writing
- MJS MacEwen: conceptualization, data collection, data analysis, validation, visualization, methodology, and writing
- J Chea: data collection, data analysis, validation, visualization
- HL Kenerson: data collection, data analysis, visualization
- AA Kwong: data collection, data analysis
- TM Locke: data collection, data analysis
- FJ Miralles: data collection, data analysis
- T Sapre: data collection, data analysis
- N Gozali: data collection
- LB Sullivan: conceptualization, data analysis
- ML Hart: data collection, data analysis
- GE Atilla-Gokcumen: conceptualization, data collection, data analysis
- S Ong: data collection, data analysis, funding acquisition, visualization
- JD Scott: conceptualization, funding acquisition
- RS Yeung: data collection, data analysis, funding acquisition
- TK Bammler: gene expression data analysis
- JW MacDonald: gene expression data analysis
- Y Sancak: conceptualization, data collection, data analysis, supervision, funding acquisition, validation, investigation, visualization, methodology, and writing

---

**Supplemental Figure 1.**
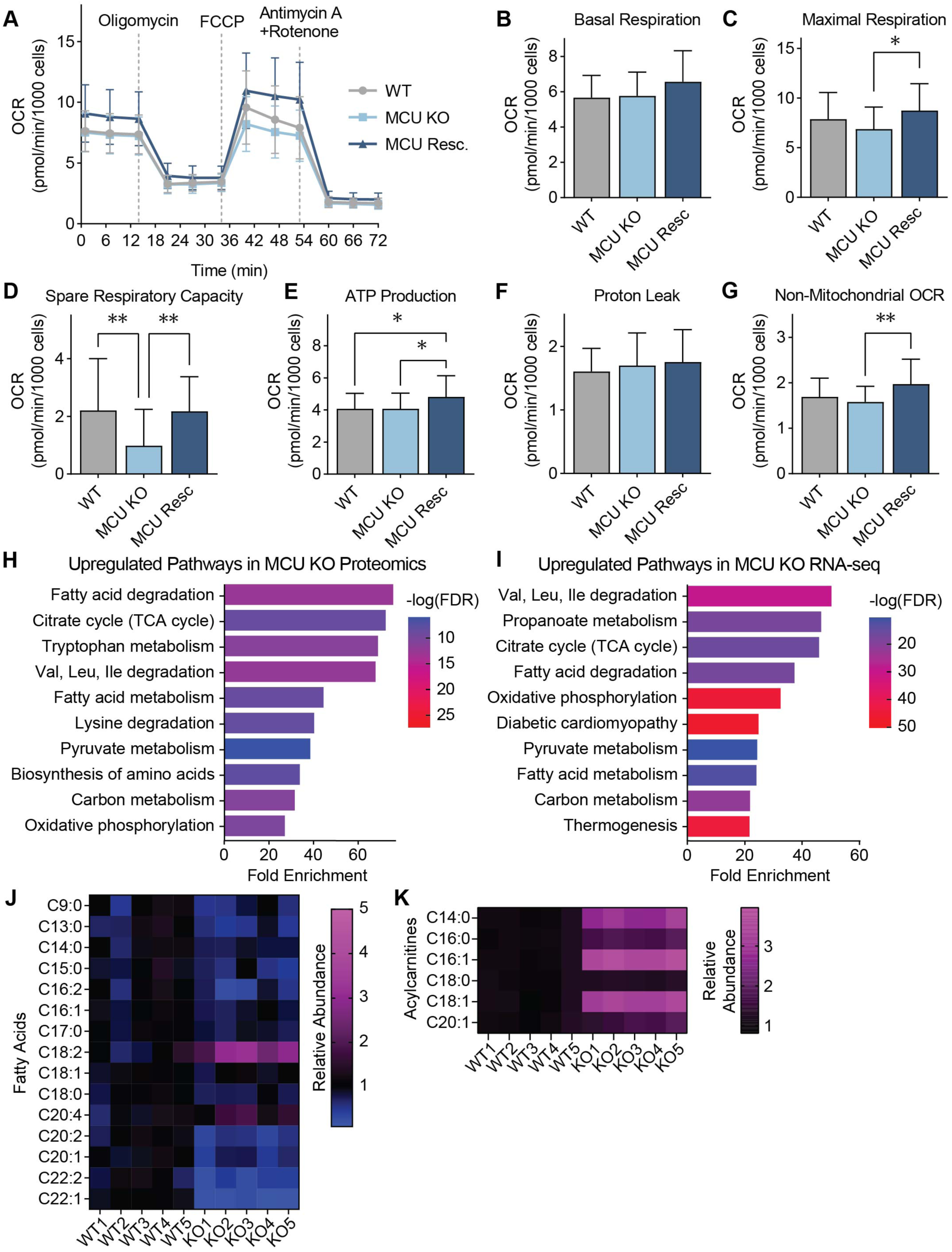
**(A-G)** Seahorse extracellular flux analysis in WT, MCU KO, and MCU rescue HeLa cells. Oxygen consumption rates at baseline and after indicated treatments are shown in (A); indicated mitochondrial parameters are shown in (B-G). Statistical significance was determined by the Tukey-Kramer test following one-way ANOVA; n=24-28. **(H, I)** Gene Set Enrichment Analysis of mitochondrial proteins (H) or mRNAs coding for mitochondrial proteins (I) that show a statistically significant increase in MCU KO cells compared to WT cells. **(J, K)** Relative abundance of fatty acids (J) and acylcarnitines (K) in WT and MCU KO HeLa cells; loss of MCU decreases steady state levels of very long chain fatty acids, but increases acylcarnitines, suggesting activation of the mitochondrial FAO pathway. All error bars indicate standard deviation; * indicates a p-value < 0.05 and ** indicates a p-value < 0.01.

**Supplemental Figure 2.**
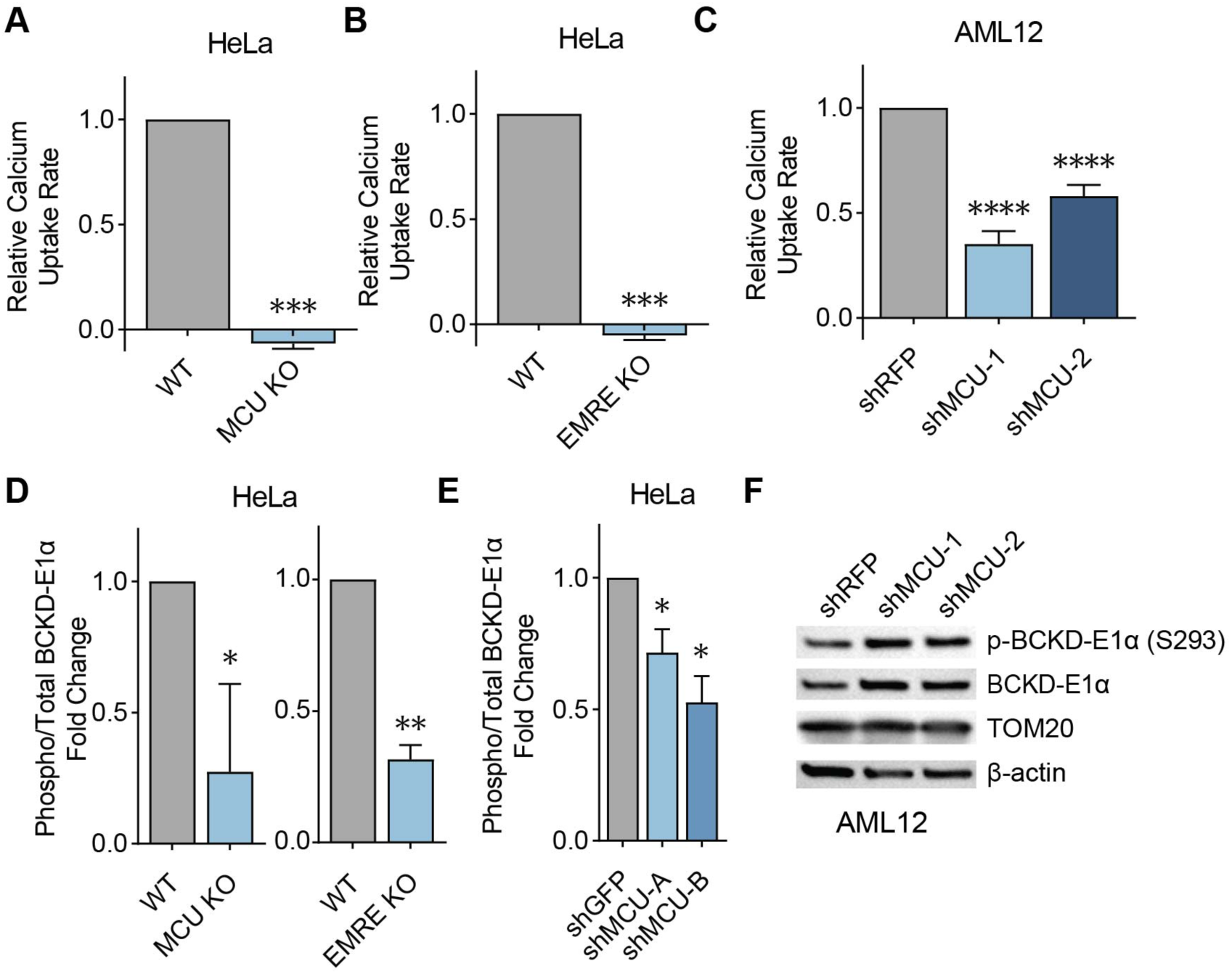
**(A,B)** Mitochondrial Ca^2+^ uptake rates in MCU KO (A) and EMRE KO (B) cells relative to WT controls are shown; n=3. **(C)** Mitochondrial Ca^2+^ uptake rates following MCU knockdown compared to control RFP knockdown in AML12 cells; n=8**. (D)** Quantification of immunoblots in Fig. 2E shown as the relative abundance of phosphorylated BCKD-E1α to total BCKD-E1α; n=3. **(E)** Quantification of immunoblots in Fig. 2F shown as the relative abundance of phosphorylated BCKD-E1α to total BCKD-E1α; n=3. **(F)** Immunoblots of phosphorylated and total BCKD-E1α in AML12 cells with or without MCU knockdown. All error bars indicate standard deviation; * indicates a p-value < 0.05, ** indicates a p-value < 0.01, *** indicates a p-value < 0.001, and **** indicates a p-value < 0.0001.

**Supplemental Figure 3.**
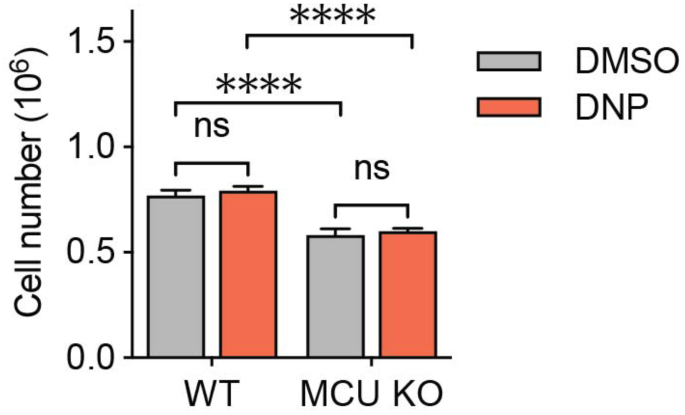
Cell numbers of WT and MCU KO cells with and without 2 μM DNP. Cells were counted three days after plating and treatment; n=3. Statistical significance was determined by Dunnett’s multiple comparisons test following one-way ANOVA

**Supplemental Figure 4.**
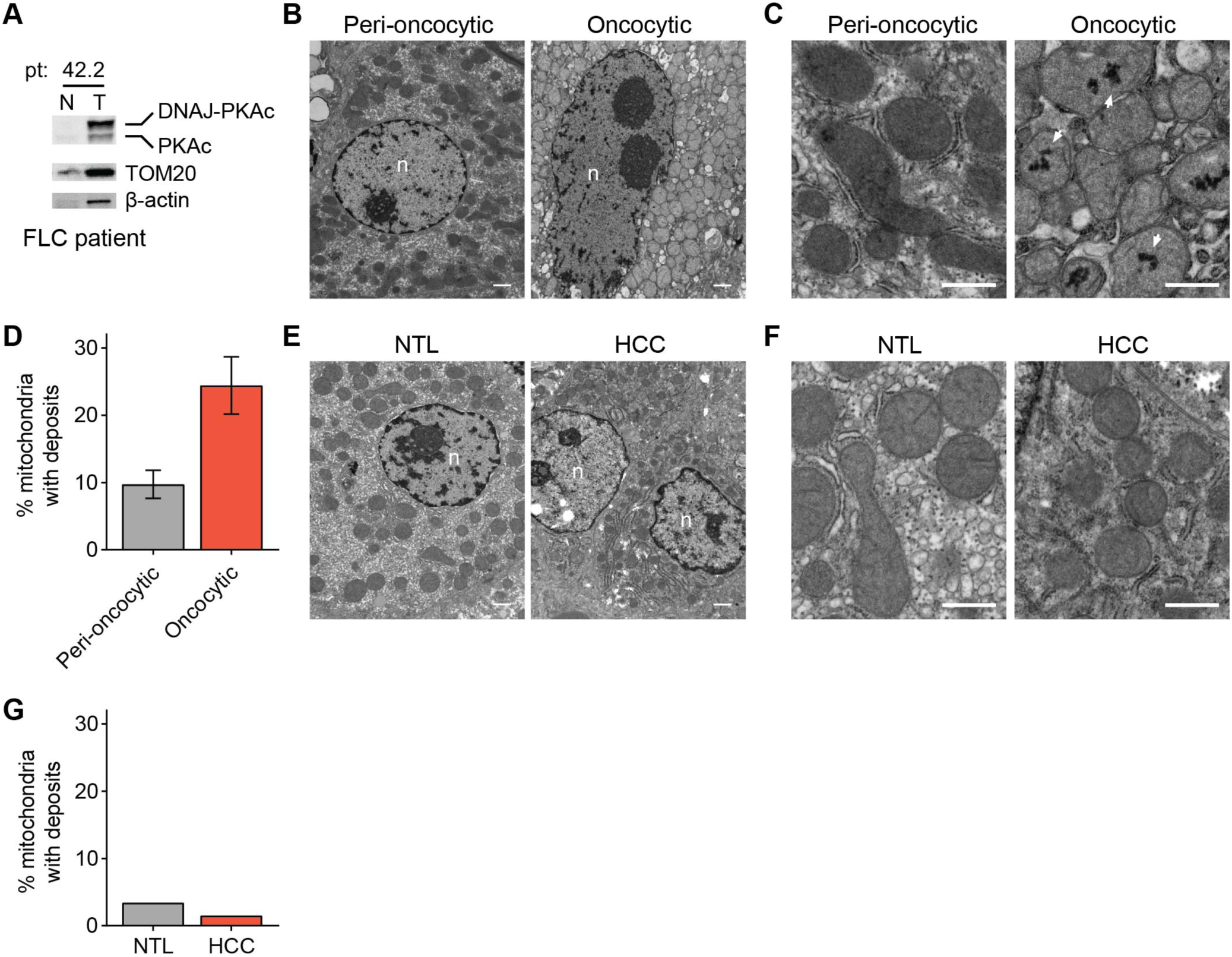
**(A)** Immunoblot of tumor (T) and non-tumor liver (N) samples from FLC patient 42.2 showing fusion protein expression. **(B)** Electron micrographs at 10,000x magnification of oncocytic liver cells and proximal (peri-oncocytic) cells from the tumor periphery of FLC Patient 42.2; normal tumor sample was not dissected in this surgery; scale bar = 1 µm; nuclei are labeled n. **(C)** Electron micrographs of samples shown in (B) at 25,000x magnification; white arrowheads mark representative Ca^2+^ deposits in the oncocytic cells; scale bar = 600 nm. **(D)** Percentage of mitochondria with Ca^2+^ deposits in EM samples shown in (C); the mean is reported from manual counting of >500 mitochondria per sample by two independent, blinded analysts; error bars indicate standard deviation. **(E)** Electron micrographs at 10,000x magnification of non-tumor (NTL) and tumor (HCC) sections from HCC patient 7; scale bar = 1 µm; nuclei are labeled n. **(F)** Electron micrographs of samples shown in (E) at 25,000x magnification; scale bar = 600 nm. **(G)** Percentage of mitochondria with Ca^2+^ deposits in EM samples shown in (F); >100 mitochondria per sample were quantified by an independent, blinded analyst.

**Supplemental Figure 5.**
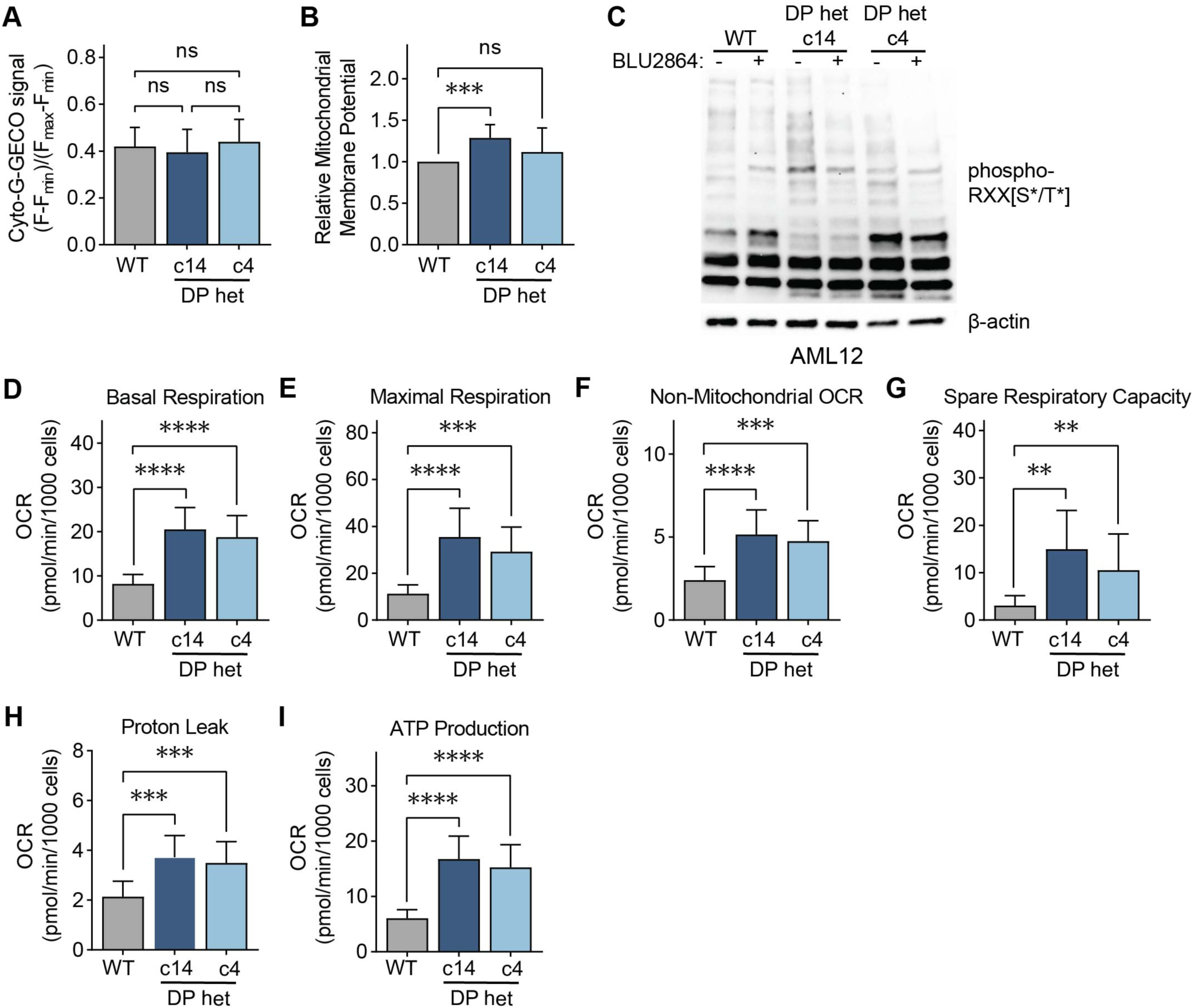
**(A)** Baseline cyto-G-GECO fluorescence normalized to minimum and maximum signals in AML12 WT, c14, and c4 cells; statistical significance was determined by one-way ANOVA; n=11-12. **(B)** Resting mitochondrial membrane potential was measured by the difference in TMRM fluorescence before and after CCCP addition, normalized to WT AML12 cells. **(C)** Immunoblot of AML12 lysates with a phospho-PKA substrate motif antibody after 5 μM BLU2864 or DMSO treatment for 4 days. **(D-I)** Indicated mitochondrial parameters of AML12 cells from Seahorse extracellular flux analysis shown in Fig. 5I; statistical significance was determined by the Dunnett test following Welch’s one-way ANOVA; n=10-16. All error bars indicate standard deviation; ns indicates non-significant, ** indicates a p-value < 0.01, *** indicates a p-value < 0.001, and **** indicates a p-value < 0.0001.

**Supplemental Figure 6.**
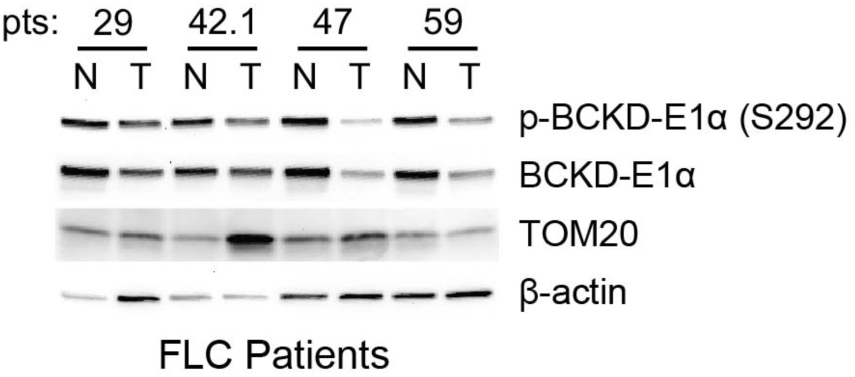
Immunoblots of phosphorylated and total BCKD-E1α in non-tumor (N) and tumor (T) lysates from FLC patients.

**Supplemental Table 1.**
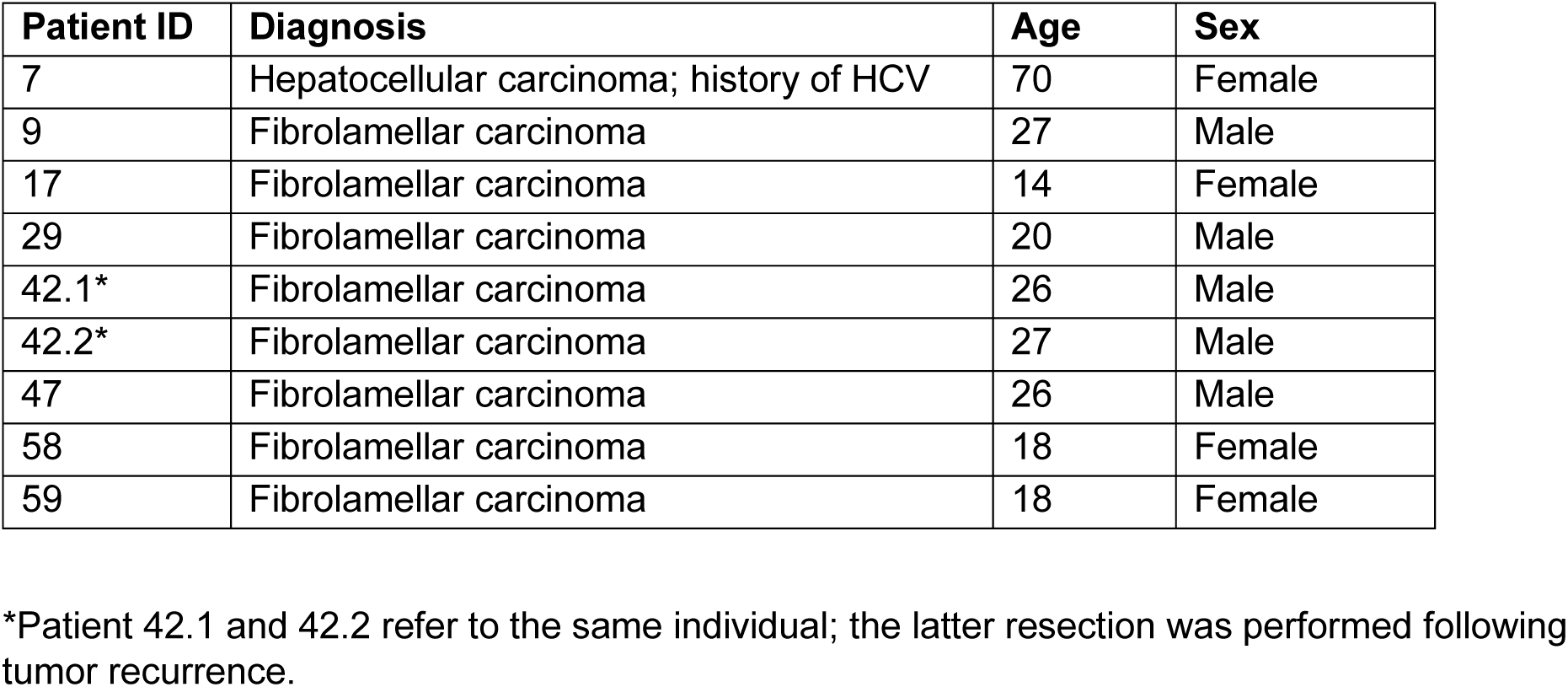
Patient Information.

| **Patient ID** | **Diagnosis** | **Age** | **Sex** |
| --- | --- | --- | --- |
| 7 | Hepatocellular carcinoma; history of HCV | 70 | Female |
| 9 | Fibrolamellar carcinoma | 27 | Male |
| 17 | Fibrolamellar carcinoma | 14 | Female |
| 29 | Fibrolamellar carcinoma | 20 | Male |
| 42.1* | Fibrolamellar carcinoma | 26 | Male |
| 42.2* | Fibrolamellar carcinoma | 27 | Male |
| 47 | Fibrolamellar carcinoma | 26 | Male |
| 58 | Fibrolamellar carcinoma | 18 | Female |
| 59 | Fibrolamellar carcinoma | 18 | Female |

\*Patient 42.1 and 42.2 refer to the same individual; the latter resection was performed following tumor recurrence.

